# Carbon usage in yellow-fleshed *Manihot esculenta* storage roots shifts from starch biosynthesis to cell wall and raffinose biosynthesis via the *myo*-inositol pathway

**DOI:** 10.1101/2023.12.07.570373

**Authors:** Sindy Gutschker, David Ruescher, Ismail Y. Rabbi, Laise Rosado-Souza, Benjamin Pommerrenig, Anna M. van Doorn, Armin Schlereth, H. Ekkehard Neuhaus, Alisdair R. Fernie, Stephan Reinert, Uwe Sonnewald, Wolfgang Zierer

## Abstract

Cassava is a crucial staple crop for smallholder farmers in tropical Asia and Sub-Saharan Africa. Although high yield remains the top priority for farmers, the significance of nutritional values has increased in cassava breeding programs. A notable negative correlation between provitamin A and starch accumulation poses a significant challenge for breeding efforts. The negative correlation between starch and carotenoid levels in conventional and genetically modified cassava plants implies the absence of a direct genomic connection between the two traits. The competition among various carbon pathways seems to account for this relationship. In this study, we conducted a thorough analysis of 49 African cassava genotypes with varying levels of starch and provitamin A. Our goal was to identify factors contributing to differential starch accumulation. With the carotenoid levels of the varieties considered as a confounding effect on starch production, we found that yellow and white-fleshed storage roots did not differ significantly in most measured components of starch or de novo fatty acid biosynthesis. However, genes and metabolites associated with myo-inositol synthesis and cell wall component production were substantially enriched in high provitamin A genotypes. These results indicate that yellow-fleshed cultivars, in comparison to their white-fleshed counterparts, direct more carbon towards the synthesis of raffinose and cell wall components, a finding that is supported by a significant rise in the starch-free residue to total dry yield ratio in yellow storage roots versus white storage roots. Our findings enhance comprehension of the biosynthesis of starch and carotenoids in the storage roots of cassava.

## Introduction

*Manihot esculenta*, also known as cassava, is a one of the most widely-cultivated crops in tropical and subtropical regions of the world, particularly in Asia and Sub-Saharan Africa, where it serves as food for millions of people (Carvalho *et al*., 2018; FAO, 2023). It is a source of carbohydrates, vitamins, and minerals, and despite the economic and nutritional importance of cassava, the yield potential of this crop has not yet been fully realized. Low productivity levels are still a major challenge faced by smallholder farmers in Sub-Saharan Africa, which leads to concerns about food security, especially in the face of climate change and a growing population. Cassava breeding programs typically prioritize the enhancement of fresh root yield and starch content due to their demonstrable influence on the adoption of new cultivars (Awotide *et al*., 2014). The most widely consumed cassava storage roots around the world are white-fleshed and starch rich, but possess low levels of micronutrients, especially provitamin A (Welsch *et al*., 2010). Vitamin A deficiency (VAD) however, still remains a prevalent problem, especially in Sub-Saharan Africa, leading not only to deterioration of vision and even blindness, but also to a weakened immune system and slow growth in children. Therefore, genetic improvement of cassava for nutritional attributes such as provitamin A are additional targets in breeding projects, intended to simultaneously address nutritional deficits (Nassar & Ortiz, 2010; Hefferon, 2015; Talsma *et al*., 2018). Initial efforts on cassava breeding aimed to enhance carotenoid or dry matter content focusing on the identification and characterization of useful genetic variants (Iglesias *et al*., 1997; Chavez *et al*., 2000). Welsch *et al*. (2010) unveiled that a C_572_A nucleotide substitution in *PHYTOENE SYNTHASE 2* (*PSY*2), a key regulator in the carotenoid biosynthesis pathway, is responsible for the qualitative color of yellow-fleshed cassava roots with additive genetic effect (AA > AC > CC). Further, the authors demonstrated that this allelic variation in *PSY*2 increases the activity of the enzyme and therefore contributes to the increased carotenoid accumulation. Subsequently, two primary loci responsible for storage root yellowness were identified by genome-wide association study (GWAS) (Rabbi *et al*., 2017). The authors also identified one yellowness locus that is co-located with a unique dry matter locus suggesting that the observed negative correlation between starch and carotenoid content might arises from physical linkage of these two loci and the allelic status of candidate genes localized within these regions (Rabbi *et al*., 2017).

Phenotypic analysis of African *Manihot esculenta* storage roots also elucidated that increased carotenoid concentrations in different cultivars correlates with reduced starch accumulation, while high carotenoid varieties still show high variations in starch accumulation levels (Njoku *et al*., 2015; Esuma *et al*., 2016; Rabbi *et al*., 2017).

Several studies examined expression or metabolite patterns in high-starch or high-carotenoid genotypes (Wilson *et al*. (2017), Beyene *et al*. (2018), Ogbonna *et al*. (2021), Xiao *et al*. (2021), Cai *et al*. (2023), Luo *et al*. (2023), Olayide *et al*. (2023)). Beyene *et al*. (2018) for example performed carotenoid enrichment research and successfully increased β-carotene content in cassava roots by coexpression of transgenes for *DEOXY-D-XYLULOSE-5-PHOSPHATE SYNTHASE (DXS)* and bacterial *PHYTOENE SYNTHASE (crtB)*, both known to play pivotal roles in carotenoid biosynthesis. This resulted in reduction of the dry matter content by 50%– 60% when compared to the non-transgenic controls and increased the concentrations of soluble sugars and triacylglycerols in the examined storage roots, especially within the ones showing the highest carotenoid levels controls. Olayide *et al*. (2023) examined the transcriptome of eleven cassava genotypes with root colors ranging from white to deep yellow, and could not observe a relationship between the expression of genes in the carotenoid biosynthesis pathway and the root’s yellowness. They hypothesized that starch and carotenoid biosynthesis might compete for glyceraldehyde-3-phosphate (GA3P). GA3P not only initiates the methyl-D-erythritol phosphate (MEP) pathway to produce precursors for ß-carotene synthesis, but is also converted to D-glucose-1-phosphate (G1P) in the starch biosynthesis pathway. However, they could not detect clear differences in expression profiles of genes which are significantly expressed in roots and involved in glycolysis between high-starch and high-carotenoid genotypes.

Despite the competition between the starch and carotenoid biosynthesis pathway, it is not yet fully understood how carbon allocation and starch accumulation in the high-carotenoid genotypes occurs, how this explains differences in carbon allocation despite the increased flux into carotenoid biosynthesis and how those strategies differ from those of high-starch/low-carotenoid varieties.

To better understand the molecular basis of the competition between starch and carotenoid accumulation, we investigated 49 field-grown African *Manihot esculenta* genotypes, with contrasting starch and carotenoid levels. We identified the transcript and metabolite patterns for high starch/low carotenoid (in the following referred to as “white” genotypes) and low starch/high carotenoid varieties (hereafter termed “yellow” genotypes) that were directly linked to the varieties starch content and showed that both groups had similar patterns in carbon allocation strategies for sucrose breakdown and starch accumulation, or *de novo* biosynthesis of fatty acids (Figure 4). However, only storage roots of yellow-fleshed genotypes partially shifted their carbon allocation away from starch accumulation and favored the accumulation of *myo*-inositol, raffinose and the biosynthesis of building blocks for pectin and hemicellulose (Figure 4). Finally, trait analyses revealed that yellow cultivars had a significantly higher ratio of starch-free residues to total dry yield (t(48) = -2.3376, p=0.022), supporting the conclusion that yellow-fleshed *Manihot esculenta* cultivars use more carbon for cell wall synthesis at the expense of their starch content (Figure 1F).

**Figure 1:**
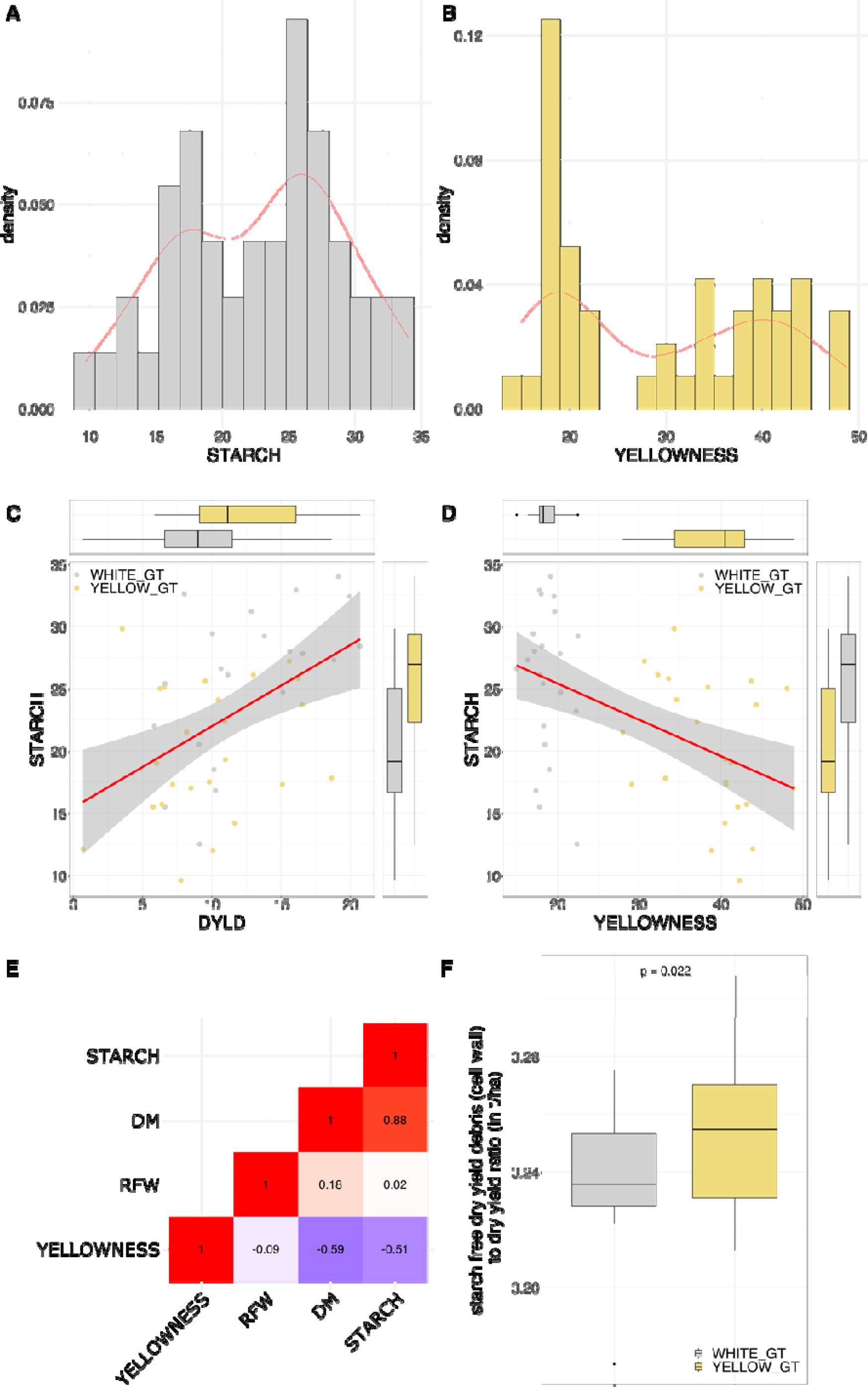
Phenotype and metabolite data representation. A-B) Density plot depicting the distribution of all genotypes under investigation according to their starch or carotenoid content. C-D) Connection between starch content (percentage) and dry yield (DYLD) or carotenoid content (yellowness; represented by chromameter b* value) and distribution across all genotypes under investigation. Yellow color refers to low starch/high carotenoid genotypes, whereas gray represents high starch/low carotenoid genotypes. Black line with the surrounding gray bar depicts the linear relationship between the variables with its corresponding 0.95 confidence interval. E) Heatmap depicting the correlation between starch, dry matter (DM), yellowness and root fresh weight (RFW). Red color indicates a positive, and blue colors a negative correlation. The filling of the heatmap boxes indicates the correlation strength. F) Depiction of difference in starch free debris to dry yield ratio between all white and yellow genotypes (both replications) under investigation and corresponding significance (t(48) = -2.3376).

## Results and Discussion

### Comparative harvest data analysis confirmed a negative correlation between starch and carotenoid content in 49 African *Manihot esculenta* genotypes

A breeding panel of 49 African *Manihot esculenta* genotypes from the International Institute of Tropical Agriculture (IITA) were grown under field condition for 12 months in Ibadan, Nigeria. Phenotype data recorded during the harvest included starch, dry matter and carotenoid content of the storage roots. As depicted in Figure 1A, starch content in the samples under investigation varied from 9-34%, depending on the genotype. Carotenoid content in cassava roots was indirectly estimated based on root yellowness intensity, as indicated by the CIELAB Chromameter b* value (Figure 1B). This method does not provide direct quantification of carotenoid levels but rather offers a relative assessment based on color intensity. The b* value, representing yellowness, ranged from 14 to 48. This color-based estimation is grounded in the known strong correlation between root yellowness and carotenoid content in cassava (Sánchez *et al*. 2014). In addition, a positive correlation between starch content and dry matter content (r ∼ 0.9, p ≤ .05) as well as a negative correlation (r ∼ -0.6, p ≤ .05) for starch and carotenoid content could be identified (Figure 1C-E). This shows that while high carotenoid varieties still accumulate starch in roots, a drop in starch levels occurs, if carotenoid content exceeds a certain amount. Besides that, it also demonstrates that yellow varieties show high variance concerning the accumulation of starch in the storage roots (Figure 1).

### Segregation of white and yellow genotypes based on metabolite profiling

Storage root samples of the 49 unique African *Manihot esculenta* genotypes from the phenotypic profiling were subjected to GC-MS analysis, and 50 metabolites were measured (Supplement material – Table S1-S2). Absolute metabolite values were z-score transformed and used to detect metabolites showing significantly different abundance between white and yellow genotypes or were correlated to starch in white or yellow varieties, respectively. In total, 16 metabolites showed a significantly different abundance between white and yellow genotypes (Figure 3). Notably, sucrose, fucose, raffinose, and xylose, along with additional metabolites, including myo-inositol, malate, fumarate, tyrosine, glutamine, and β-alanine exhibited statistically significant decrease in abundance in white relative to yellow genotypes (Figure 3). Only three metabolites - nicotinic acid, 4-hydroxy-*trans*-proline, and glutamate - displayed significantly higher abundance in the white compared to the yellow genotypes. When analyzing the correlations between starch content and the metabolites showing significant different abundance between the two genotype groups, only 4-hydroxy-*trans*-proline revealed a significant positive correlation with starch in the yellow genotypes. The remainder were consistently negative in both white and yellow genotypes (Figure 3).

The differences in metabolite abundance, especially the higher abundance of sucrose, glucose and fructose, indicate that it is unlikely that the reduced starch formation in yellow genotypes is due to limited source supply of carbon. Sucrose, fructose and glucose were also negatively correlated with starch in both white and yellow genotypes, suggesting a differentiation in carbon allocation rather than reduced photosynthetic performance in the yellow varieties. This accumulation of sugars in the storage root of low dry matter genotypes has also been reported in a smaller subset of the same African cassava genotype panel by Lamm *et al*. (2023), recently.

### Segregation into white and yellow genotypes based on *PSY2* sequence

The same storage root samples of the 49 unique African *Manihot esculenta* genotypes were subjected to RNA sequencing using the protocol of the Illumina NovaSeq6000 system (PE150+150; Novogene, Cambridge, UK). After removal of adapters and quality trimming (Q35), an average of 21 million paired-end reads per sample were retained. Sequence alignment to the reference genome yielded an average mapping efficiency of 91% for uniquely mapped reads and 4% for multiply mapped reads.

Sequence analyses of the *PSY*2 reads showed that the genotype groups carry different alleles for that locus. All varieties characterized as “yellow”, showed the same A to C substitution in the *PSY*2 transcript (Chromosome 1, Position 31960252 in *Manihot esculenta* v8.1 reference genome) as described by Welsch *et al*. (2010). All white genotypes showed the null version of it. This not only supports the distinction into white and yellow genotypes, but also demonstrates that carotenoid biosynthesis in yellow-fleshed *Manihot esculenta* storage roots might be mainly driven by the allelic differences in *PSY*2 (Supplementary material – Table S3). However, it is unclear which regulations drive the differential starch accumulation in yellow genotypes.

### Transcriptional separation and correlation analysis for white and yellow varieties

All uniquely mapped reads were additionally subjected to variance-stabilizing transformation and depth filtration (1710), identifying approximately 20.000 genes as being expressed at least in one of the genotype groups. Principal component analysis (PCA) of all expressed genes revealed distinct clustering patterns distinguishing white from yellow genotypes along PC1. In particular, the lack of an overlap between the clusters for white and yellow genotypes underscores a transcriptional divergence that accounts for 61% of the observed variance in PC1 (Figure 2A). A follow-up differential expression analysis uncovered a total of 472 differentially expressed genes (DEGs) between white and yellow genotypes with an even distribution across all chromosomes (Figure 2B-C and Supplementary material – Figure S1). Of these, 71 DEGs were higher expressed in white and 401 in yellow varieties (Figure 2B – and Supplementary material – Table S4). However, comparable to the results of Olayide *et al*. (2023), almost no significant difference in expression of genes being annotated to be part of the carotenoid biosynthesis pathway could be detected between white and yellow genotypes. This is in agreement with the finding that the C to A substitution in *PSY2* increases its enzyme activity (Welsch *et al*., 2010) and suggests that differences in carotenoid biosynthesis in the yellow-fleshed *Manihot esculenta* storage roots analyzed here are mainly driven by the allelic differences in *PSY*2. Only *DXS*, catalyzing the first step in the MEP pathway, was detected with a significantly higher abundance in yellow compared to white genotypes (Figure 4 and Table 2). In *Solanum lycopersicum* and *Arabidopsis thaliana*, an enhanced enzyme activity of PSY could be observed together with a post transcriptional accumulation of DXS (Fraser *et al*., 2007; Rodríguez-Villalón *et al*., 2009). Studies in Arabidopsis have shown that DXS is the enzyme with the highest flux control through the carotenoid biosynthesis pathway (Wright *et al*., 2014). This might explain the higher transcript levels of *DXS* in yellow *Manihot esculenta* storage roots, although the reported allele-specific enzyme activity increase of *PSY2* (Welsch *et al*., 2010) likely has a bigger effect on carotenoid levels, also in the genotypes analyzed here.

**Figure 2:**
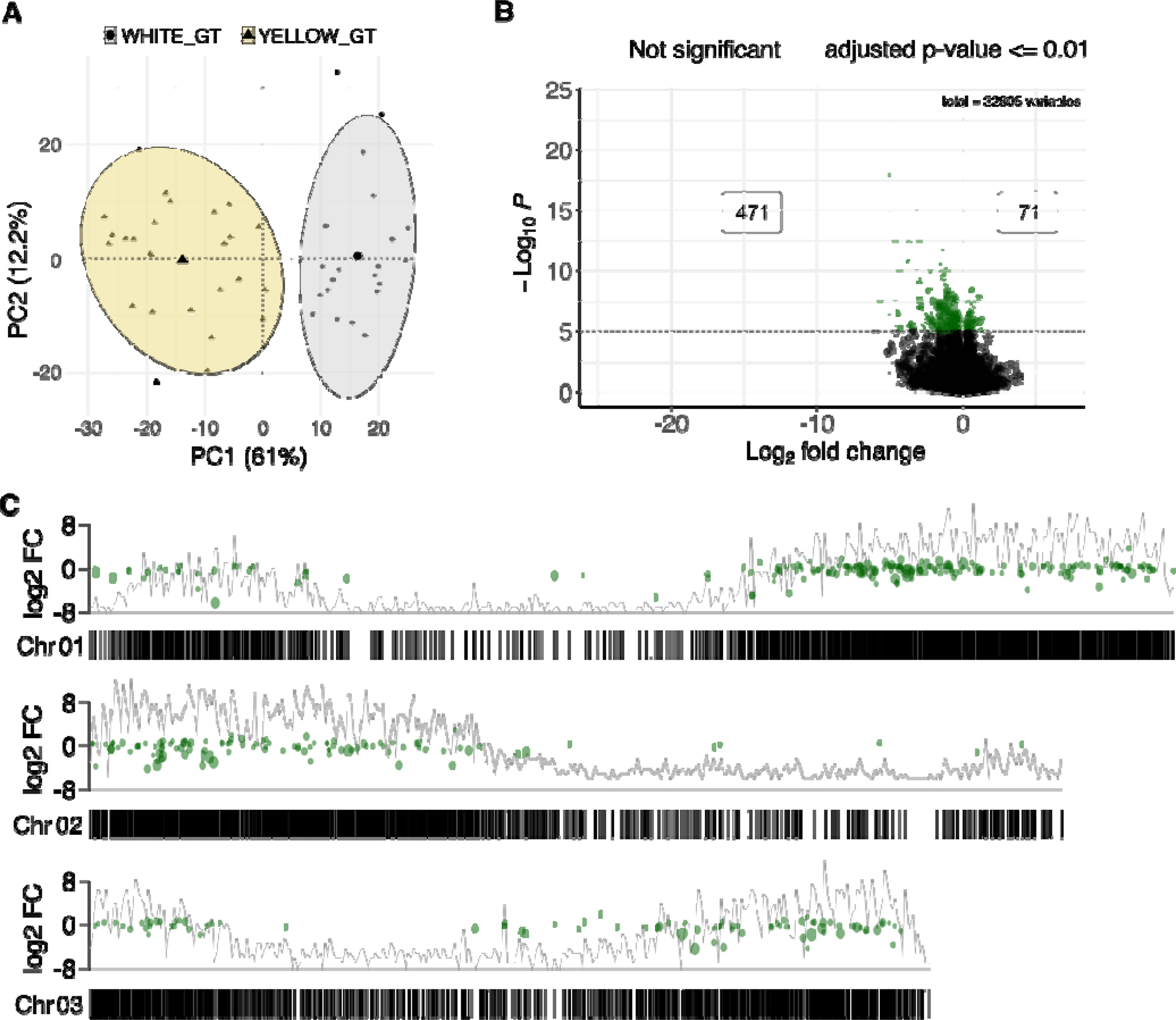
Representation of transcriptional information and differences between examined genotype groups. A) Principal component analysis representing PC1 and PC2 for read count-filtered transcriptomic data. Small triangles and yellow circle label yellow genotypes, whereas the gray circle and small black dots represent white genotypes. Enlarged triangle and black dot highlight the center of the corresponding genotype distribution in PC1 and PC2 dimensions. B) Volcano plot representing all genes between white and yellow genotypes. Black dots depict genes with no significant difference between white and yellow genotypes. Green dots refer to genes that match the significance threshold. Positive Log2FC values refer to a higher expression in white, whereas negative values represent a higher expression in yellow genotypes. C) Chromosomal distribution of all detected significantly differential expressed genes between white and yellow genotypes. Black bars at the bottom depict the localization of genes on the respective chromosome. The line in the background represents the number of transcripts for that position. Green dots refer to the significant differentially expressed genes for that chromosome. Y-axis and equivalent height of green dots indicate changes in gene expression comparing white with yellow genotypes – positive values refer to a higher expression in white, whereas negative values representing a higher expression in yellow genotypes. For conciseness, only the first three chromosomes are shown. For representation of all chromosomes see Supplementary material – Figure S1.

Our analyses revealed not only a negative correlation between starch and carotenoid content and an opposite trend in white and yellow genotypes. We also show that yellow cultivars have different starch contents despite hetero- or homozygosity for the C_572_A nucleotide substitution in *PSY2* and the resulting characterization as yellow-fleshed. Therefore, the question arises on how carbon is allocated within the yellow genotypes, how the variance in starch accumulation despite the carotenoid levels can be explained and compared to the mechanisms in white genotypes.

Considering the negative correlation between starch and carotenoid content, direct analyses between gene expression and starch content without accounting for carotenoid levels could result in misleading conclusions. Specifically, gene expressions associated with starch metabolism might unintentionally reflect the influence of the correlation between starch and carotenoid levels. To precisely capture the true relationships between gene expression and starch content solely, we performed a partial correlation analysis. This statistical approach enabled us to capture the relationship between gene expression and starch content and to filter out potential (direct or indirect) influences from the variety’s carotenoid content. Therefore, high-quality, depth-filtered (1710) reads that underwent variance-stabilizing transformation were used to identify genes with expression profiles significantly correlated to starch content, while accounting for the variety’s carotenoid levels as a confounding effect performing partial correlation, separately for white and yellow genotypes. Using this approach, as represented in Table 1, white genotypes showed more genes being significantly correlated to starch in total, while at the same time a proportionally higher number of them being positively correlated to starch (see Supplementary material – Table S5). Functional enrichment analysis for the white genotypes revealed, that the expression of positively correlated genes were enriched in the “Protein processing in endoplasmatic reticulum”, “Nucleotide metabolism”, “Pyrimidine metabolism”, and “Thiamine metabolism” Kyoto Encyclopedia of Genes and Genomes (KEGG) pathways. Yellow varieties, on the other hand, co-increase, among others, genes related to “Starch and sucrose metabolism”, “Amino sugar and nucleotide sugar metabolism,” and “Biosynthesis of nucleotide sugars”. For genes negatively correlated to starch content within the white and yellow genotypes, an enrichment of genes annotated as part of “Ribosomes”, “D-Amino acid metabolism” and “Ribosome biogenesis in eukaryotes” could be detected (see Supplementary material - Figure S2).

**Table 1:** Number of genes correlated to starch, separated for white and yellow genotypes.

| Number of genes correlated to starch | White genotypes | Yellow genotypes |
| --- | --- | --- |
| Positive correlation | 1024 | 665 |
| Negative correlation | 942 | 736 |
| $\Sigma$ | 1966 | 1401 |

### White and yellow genotypes show positive correlation of genes involved in starch biosynthesis

In *Manihot esculenta,* photoassimilates produced in photosynthetically active leaves (source tissue) are converted to sucrose, transported to the roots via the phloem tissue and vascular rays and symplasmically to the xylem parenchyma of storage roots (sink tissue) (Mehdi *et al*., 2019). As depicted in Figure 4, incoming sucrose can then be cleaved into glucose and fructose by CYTOSOLIC INVERTASES (cINV) and further phosphorylated by HEXO- or FRUCTOKINASES (HXK; FRK) to form glucose- or fructose-6-phosphate (G6P; F6P). GLUCOSE-6-PHOSPHATE ISOMERASE can act as an intermediate catalyst for the conversion of F6P to G6P. However, *Manihot esculenta* mainly utilizes SUCROSE SYNTHASE (SUS) in its storage roots to cleave incoming sucrose (Mehdi *et al*., 2019) into UDP-glucose and fructose, the latter of which is phosphorylated by FRK, while UDP-GLUCOSE PYROPHOSPHORYLASES (UGPases) and PHOSPHOGLUCOMUTASES (PGMs) catalyze the reaction of UDP-glucose to G1P and then to G6P, respectively. When these genes were analyzed in terms of expression patterns between white and yellow genotypes and their correlation to starch while accounting for carotenoid levels as a confounding effect (Figure 4, Table 3 and Supplementary material – Table S5-S6), only one of three *cINV*s was detected as being higher expressed in white compared to yellow genotypes. The remaining two isoforms were not differentially expressed between white and yellow genotypes. In addition, none of them showed an expression pattern correlated with starch in either genotype group, even though all *cINV*s annotated transcripts were expressed in both genotype groups. Also, *SUS* did not show significant differential expression between the genotype groups, but one isoform was positively correlated to starch in white genotypes. Furthermore, six out of nine *SUS* annotated transcripts were similarly expressed in both genotype groups. One *UGPase* was detected with significantly higher abundance in yellow genotypes, while a different isoform was positively correlated to starch in white genotypes. The two cytosolic isoforms of *PGM* did not show any significant correlation to starch or differential expression between the genotype groups but were expressed in both white and yellow genotypes (Figure 4, Table 3 and Supplementary material – Table S5-S6).

**Table 2:** Enzymes related to carotenoid biosynthesis and identified as correlated to starch or significant differentially expressed between white and yellow genotypes.

| Enzyme | Isoform | Locus identifier | Corr. coeff<br>WHITE | Corr. coeff<br>YELLOW | Expression sign.<br>higher in |
| --- | --- | --- | --- | --- | --- |
| 1-deoxy-D-xylulose 5-phosphate synthase | DXS1 | Manes.15G018800 | n.s. | n.s. | YELLOW |
|  | DXS1 | Manes.13G101600 | n.s. | n.s. | YELLOW |
| 4-cytidine-5'-diphospho-2-C-methyl-D-erythritol kinase | CMK | Manes.01G176400 | n.s. | -0.45 | n.s. |
| Phytoene synthase | PSY | Manes.02G081700 | 0.70 | n.s. | n.s. |

**Table 3:**
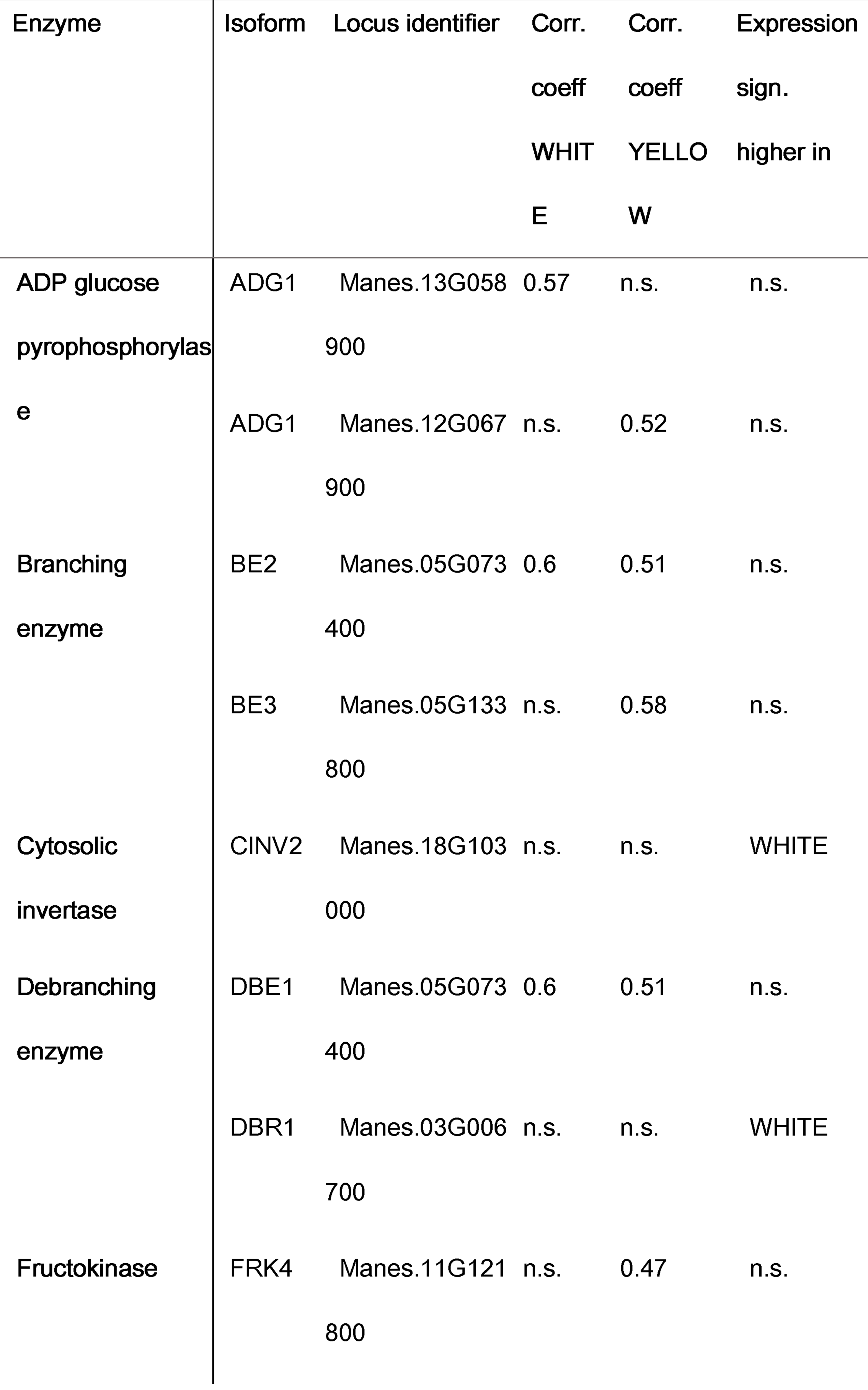
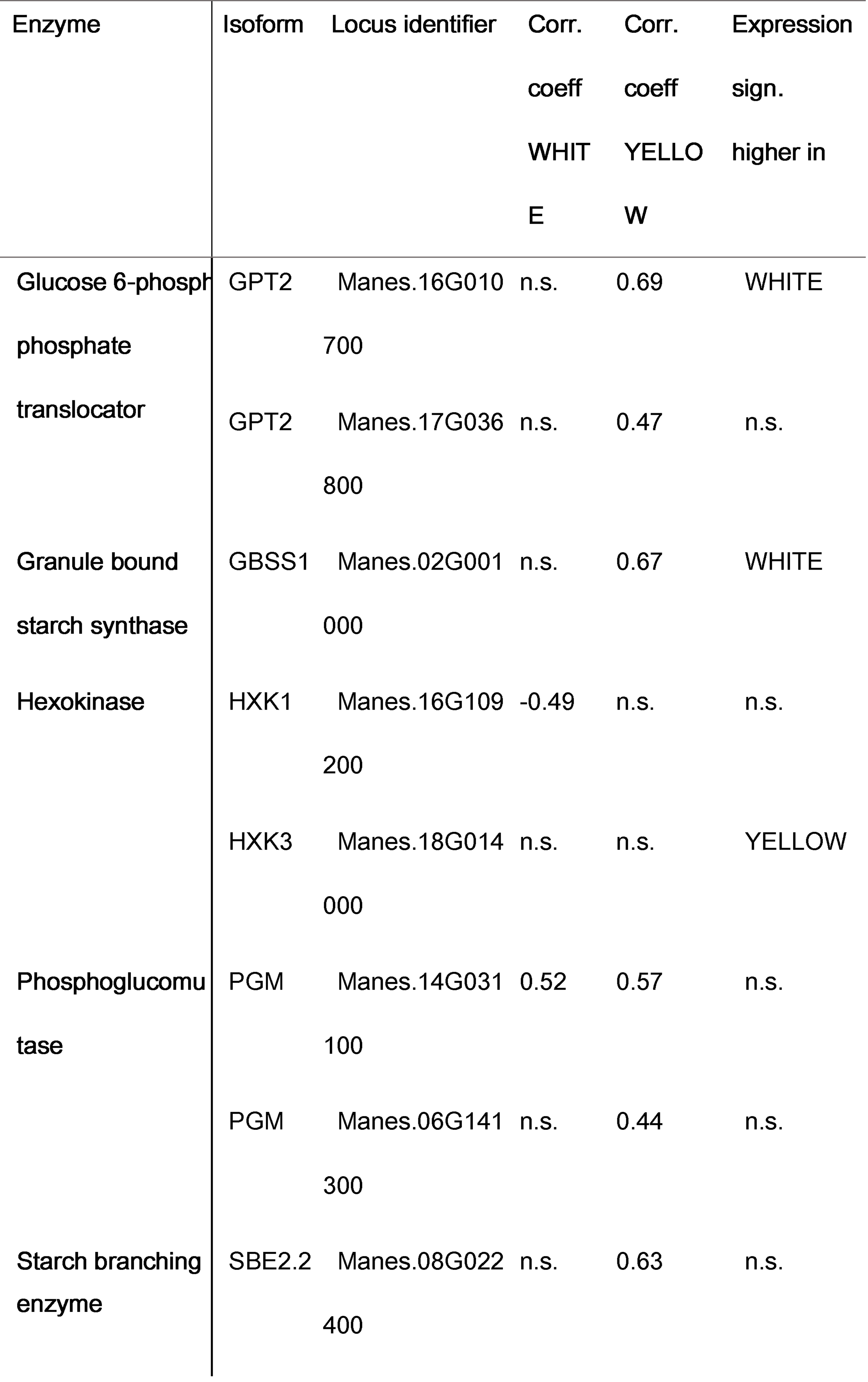
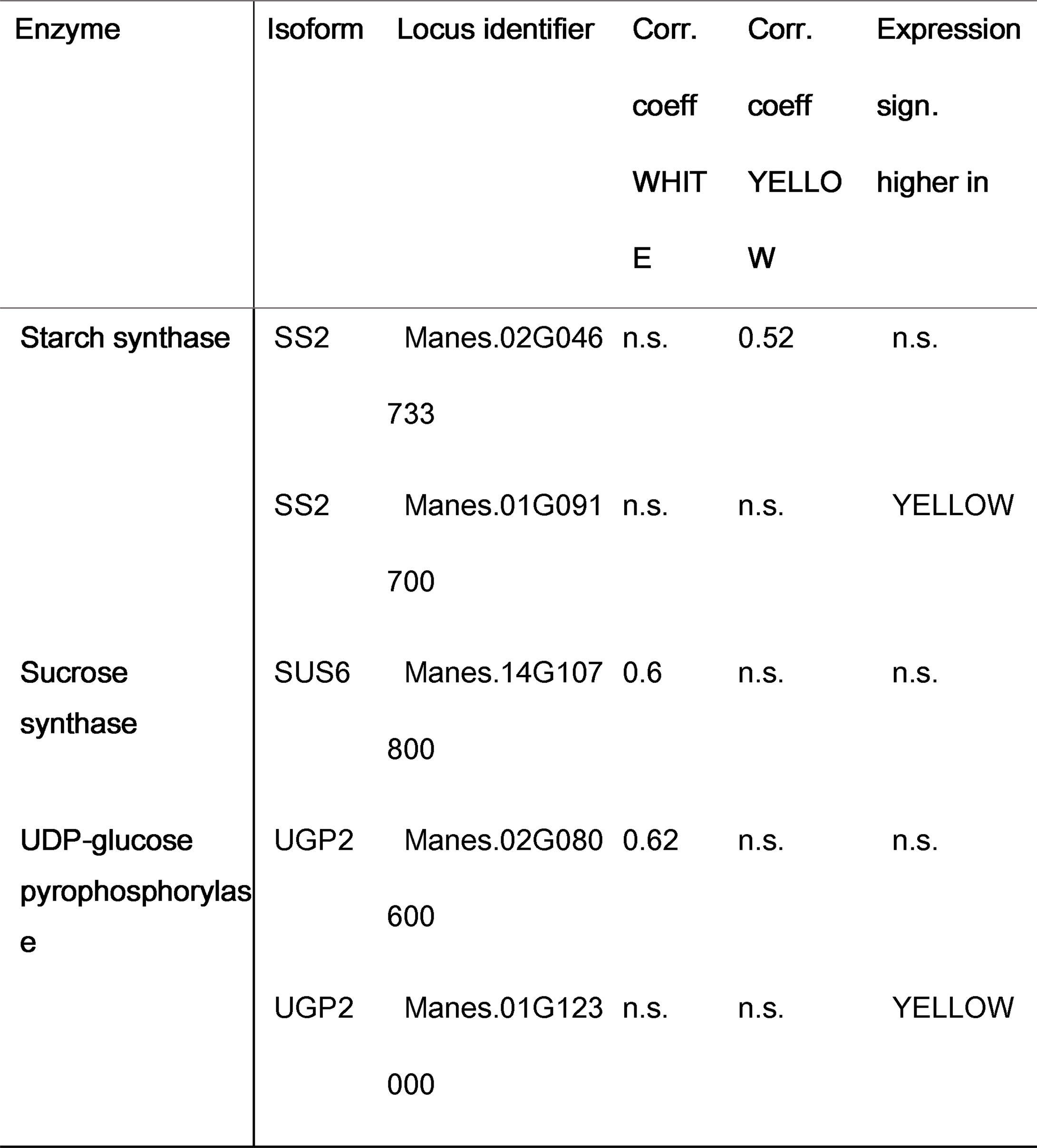
Enzymes related to starch biosynthesis and identified as correlated to starch or significant differentially expressed between white and yellow genotypes.

In conclusion, although we were able to detect one *SUS* isoform (Table 3) that positively correlated to starch in the white genotypes, we were unable to detect differential expression between the genotype groups studied for *SUS*. For *cINVs* only one isoform appeared differentially expressed between the white and yellow genotypes analyzed here. Despite a higher sucrose content, higher glucose and fructose levels were also observed, albeit not significantly, in the yellow cultivars, which may indicate a lower flux through the SUS-mediated metabolic pathway and a preference for the cINV metabolic pathway in the yellow cultivars studied here. Similar observations were also made by Lamm *et al*. (2023), who investigated metabolic and proteomic differences in eight *Manihot esculenta* genotypes of the same genotype panel. The authors found a shift from cINV-to SUS-mediated sucrose cleavage and an increased allocation of sugars to the amyloplasts in high dry matter genotypes.

G6P can enter the amyloplast via the GLUCOSE-6-PHOSPHATE/PHOSPHATE TRANSLOCATOR (GPT) and is converted back to G1P by PGMp. G1P then enters the rate-limiting step, conversion to ADP-glucose, catalyzed by ADP-GLUCOSE PYROPHOSPHORYLASE (AGPase), providing the building blocks for starch. In the genotypes analyzed here, one of two *GPT2* isoforms was identified as DEG between white and yellow genotypes, with higher expression in white genotypes and both isoforms were detected as positively correlated to starch only in yellow genotypes. This specific difference between white and yellow genotypes suggests that the uptake of G6P into the amyloplasts might be an important point of metabolic divergence between white and yellow genotypes. The importance of carbon and energy uptake into amyloplasts for starch synthesis was also demonstrated in potato tubers, previously (Jonik *et* al., 2012, Zhang *et al*., 2008). The two plastidial *PGM* isoforms showed the same expression levels between the genotype groups. One with positive correlation to starch in both groups, the second with positive correlation only in yellow genotypes. Four *AGPase* subunits were identified with no differential expression comparing white and yellow genotypes. However, for both genotype groups a positive correlation between the expression of *AGPase* and starch could be observed, although a different isoform of *AGPase* significantly correlated in each genotype group.

Finally, amylopectin and amylose, the two main components of starch, are produced for storage by GRANULE BOUND STARCH SYNTHASE (GBSS), STARCH SYNTHASE (SS) and STARCH BRANCHING ENZYME (BE) (Munyikwa et al., 1997; Figueroa *et al*., 2022). Furthermore, DEBRANCHING ENZYMES (DBE) are known to partake functions in amylopectin synthesis as well, even though it was confirmed that those are not strictly necessary for starch formation (Zeeman *et al*., 2010). One *GBSS* isoform showed a higher abundance in white genotypes and a positive correlation to starch in the yellow genotypes examined here. One *SS* and two *BE* isoforms exhibited a positive correlation to starch in yellow genotypes and one *SS* isoform additionally a higher expression in yellow genotypes (Figure 4, Table 3 and Supplementary material – Table S5-S6).

All in all, both genotype groups share similar gene expression and correlation patterns for genes known to be involved in sucrose breakdown and starch biosynthesis.

### White and yellow genotypes show only minor differences in expression of genes involved in *de novo* fatty acid biosynthesis and their correlation to starch

Increased carotenoid levels in *Manihot esculenta* co-occur not only with an decrease in starch content, but were also shown to coincide with an increase in fatty acid concentrations (Beyene *et al*. 2018) which indicates a competition for carbon skeletons between *de novo* fatty acid and starch synthesis due to redirection of carbon resources.

To address whether there is a transcriptional or metabolic basis for these observations in the genotypes analyzed here, we investigated the gene expression and metabolite abundance known to partake in *de novo* fatty acid biosynthesis (Guschina & Harwood, 2007; Harwood, 1996; Figure 4 and Table 4 and Supplementary material – Table S5-S6). White and yellow genotypes only exhibit small alterations in gene expression associated with *de novo* fatty acid synthesis and correlations to starch were almost exclusively negative. As depicted in Figure 4 and Table 4, one *3-KETOACYL-ACP SYNTHASE (KASII)* isoform was detected to be negatively correlated to starch and significantly higher abundant in white genotypes. *KASI* was identified as being higher expressed in white genotypes but did not show any correlation to starch and one *ß-KETOACETYL-CoA REDUCTASE* (*KCR)* isoform was identified with a negative correlation to starch in white genotypes. One *FATTY ACYL-ACP THIOESTERASE (FaTB)* isoform was examined with a positive correlation to starch, while a second one was higher expressed in the yellow compared to the white genotypes. None of the other genes involved in *de novo* fatty acid synthesis showed significant differential expression between white and yellow genotypes, even though almost all of them were detected as expressed in both genotype groups (Figure 4 and Table 4 and Supplementary material – Table S5-S6). In addition, acetyl-CoA may be proposed as a competing metabolite between starch and fatty acid synthesis as well. Contributing to *de novo* fatty acid synthesis since providing building blocks for malonyl-CoA, acetyl-CoA can be synthesized (1) from free acetate via the action of ACETYL-CoA SYNTHETASE (ACS), (2) from pyruvate by the reaction catalyzed by PYRUVATE DEHYDROGENASE COMPLEX (PDC) in the plastid or mitochondria, (3) from free acetate through the action of a plastidic CARNITINE ACETYLTRANSFERASE which transfers acetate from acetyl-carnitine to CoA or (4) through the ATP-CITRATE LYASE (ACL) (Ke *et al*., 2000; Rawsthorne, 2002). Nonetheless, we could not detect differential expression between the white and yellow genotypes, nor a significant correlation to starch of aforementioned acetyl-CoA synthesizing genes, or additional genes taking place in the TCA cycle or glycolysis, with the exception of one *PHOSPHOGLYCERATE KINASE* (*PGK*3) isoform being significantly higher abundant in white genotypes and positive correlation to starch in yellow varieties and two *E1 ALPHA SUBUNIT OF PYRUVATE DEHYDROGENASE COMPLEXES (PDC’s)* being significantly higher expressed in yellow genotypes, with one of them positively correlated to starch in white varieties (Supplementary material Table S4-S6).

**Table 4:**
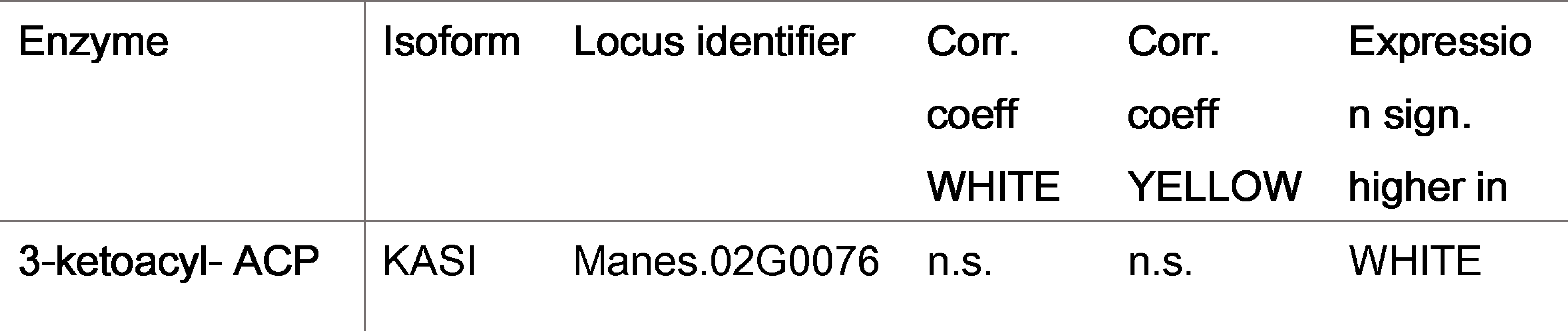
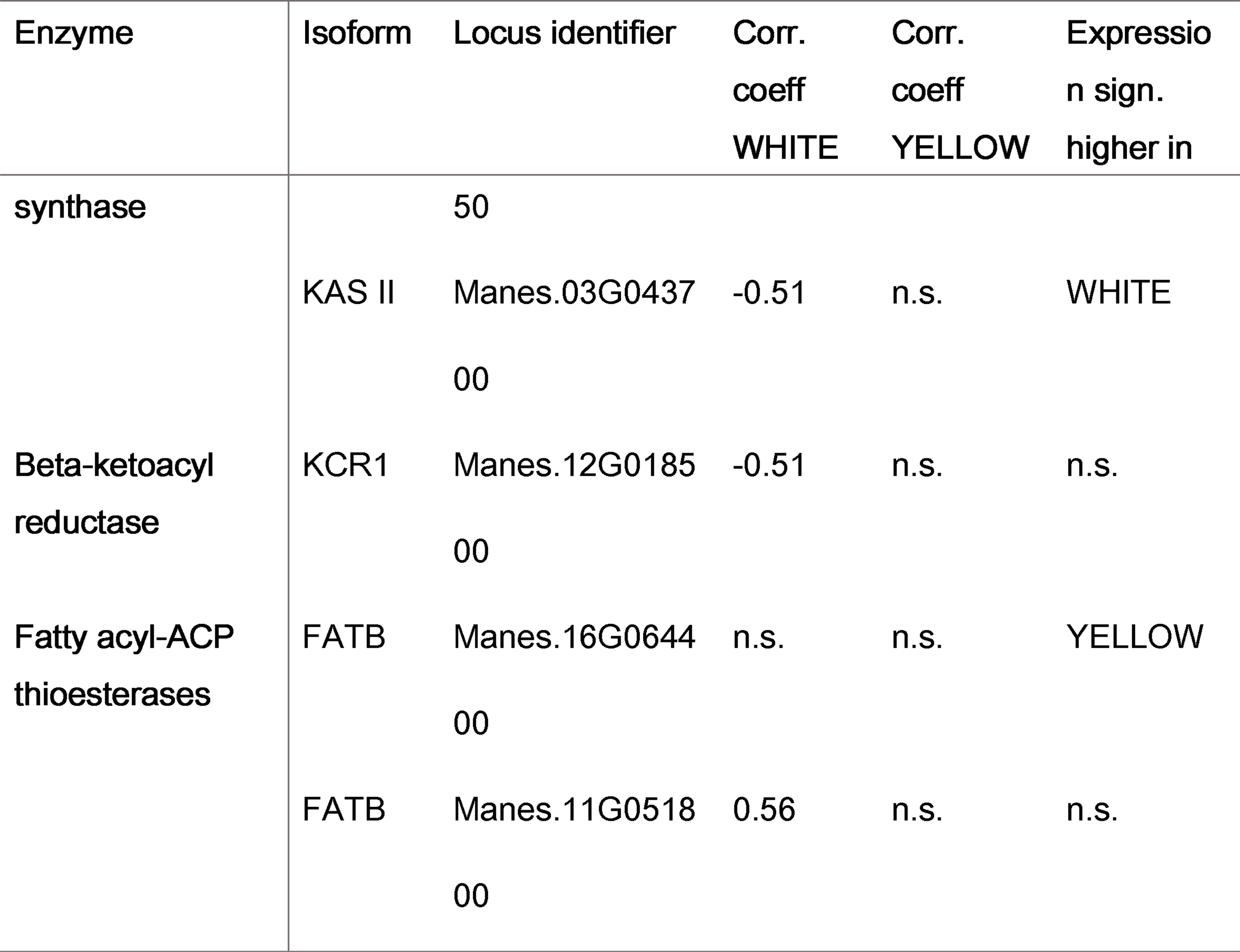
Enzymes related to the *de novo* fatty acid biosynthesis and identified as correlated to starch or significant differentially expressed between white and yellow genotypes.

Based on these results, it appears that the transcriptome data and gene expression patterns in the *Manihot esculenta* genotypes studied, do not convincingly support a clear shift from starch to fatty acid synthesis in the yellow varieties. While it was observed that lower starch levels and higher carotenoid concentrations co-occurred with increased fatty acid concentrations (Beyene et al., 2018), the transcriptional and metabolic evidence suggests that starch and fatty acid synthesis may not necessarily directly compete with each other in the varieties analyzed here. This observation is in line with previous research in other plant species - e.g. Klaus et al. (2004), who analyzed among others, potato lines with reduced activities of AGPase or plastidial PGM to test re-direction of carbon to fatty acid synthesis and revealed that the block in the limiting steps in starch synthesis did not result in a redirection of carbon into lipid synthesis in *Solanum tuberosum* L. plants. All in all, despite some alterations in the expression of certain genes related to de novo fatty acid biosynthesis, the overall lack of significant differential expression for most of these genes imply a complex metabolic regulation that allows both pathways to coexist but does not point towards a clear and convincing direct trade-off.

### Transcript and metabolite levels highlight significant higher abundance of genes and metabolites associated with *myo*-inositol, raffinose and cell wall biosynthesis in yellow genotypes

*Myo*-inositol in plants serves many diverse functions. To name just a few, it is a key metabolite for signaling and cellular communication, a precursor for phosphate storage (in form of phytate), and is needed for cell wall biosynthesis and the production of stress related oligosaccharides like raffinose or stachyose.

*Myo*-inositol can be synthesized *de novo* from G6P by converting it to *myo*-inositol-1-phosphate by the action of *MYO*-INOSITOL-1-PHOSPHATE SYNTHASE (MIPS) in the rate limiting step of inositol biosynthesis and to *myo*-inositol by the reaction catalyzed by *MYO*-INOSITOL PHOSPHATASES (IMP).

In cell wall biosynthesis, UDP-glucuronic acid is synthesized via two different pathways. The primary pathway involves the direct conversion of UDP-glucose to UDP-glucuronic acid by UDP-GLUCOSE DEHYDROGENASE (UGD). In parallel, the secondary pathway involves the *myo-*inositol pathway. *Myo*-inositol serves as a building block for glucuronic acid formed via *MYO*-INOSITOL OXYGENASE (MIOX) which in turn leads to the synthesis of UDP-glucuronic acid (UDP-GlcA) via GLUCURONOKINASES (GLCAK) and UDP-SUGAR PYROPHOSPHORYLASE (USP). By the action of UDP-D-APIOSE/UDP-D-XYLOSE SYNTHASE (AXS), UDP-XYLOSE SYNTHASE (UXS), UDP-D-XYLOSE 4-EPIMERASE (UXE), and UDP-D-GLUCURONATE 4-EPIMERASE (GAE). Subsequently, UDP-GlcA will be used to form UDP-arabinose, UDP-apiose, UDP-galacturonic acid, and UDP-xylose, all of which are important cell wall components and building blocks for pectin or hemicellulose (Figure 4). As depicted in Figure 3 and 4, sucrose, *myo*-inositol, and xylose were significantly higher in yellow compared to white genotypes analyzed here. In addition, *MIPS*, *USP* and *UXS* were detected with a significantly higher expression in yellow compared to white genotypes (Figure 4, Table 5 and Supplementary material – Table S5-S6). Furthermore, *MIPS*, *UXE* and *GAE* were identified with a significant negative correlation to starch in the yellow genotypes. Only *UXS* was detcted with a negative correlation and only *UXE* with a positive correlation to starch in the white genotypes. Only *AXS* showed a positive correlation to starch in the yellow genotypes in the pathways described above (Figure 4, Table 5 and Supplementary material Table S5-S6). Collectively, yellow varieties seem to not significantly prefer the UGD pathway for glucuronic acid synthesis. However, the significantly higher *MIPS* abundance, it’s negative correlation to starch, and the significantly higher abundance of *myo*-inositol together with a higher expression of *USP* in yellow genotypes supports the hypothesis of a preferred UDP-D-Glucuronic acid synthesis via the *myo*-inositol pathway. The detected negative correlation of UXE expression and starch, the negative correlation to starch of *GAE* and the higher abundance of xylose in the yellow varieties studied here (Figure 3) further suggest that a re-direction of sucrose or G6P to the *myo*-inositol pathway in yellow-fleshed genotypes lead to an increased synthesis of pectin and hemicellulose building blocks.

**Figure 3:**
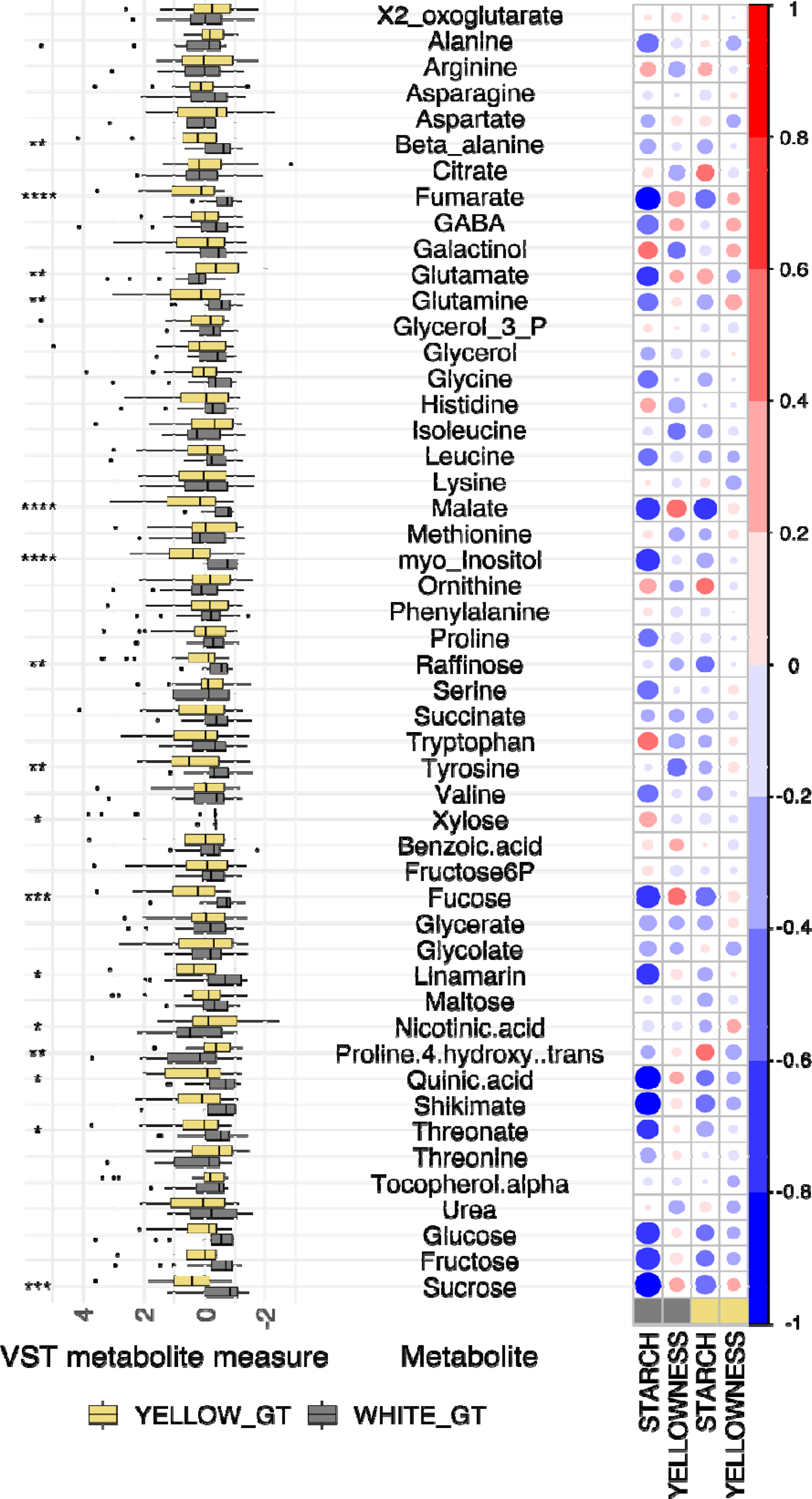
Representation of metabolite information, differences between examined genotype groups and their correlation to starch and carotenoid content. The left side represents all measured metabolites and their difference in abundance between white and yellow genotypes. Yellow color refers to low starch/high carotenoid genotypes, whereas gray represents high starch/low carotenoid genotypes. The right side depicts correlation analysis of all measured metabolites to starch and yellowness. Boxes at the bottom represent low starch/high carotenoid (yellow) and high starch/low carotenoid (grey) genotypes. Red colors indicate a positive, and blue colors a negative correlation. The circle sizes display the correlation strength. Asterisks (*; ** and ***) are refer to the significance level of the corresponding p-value (≤.05; ≤.01 and ≤.001)

**Table 5:**
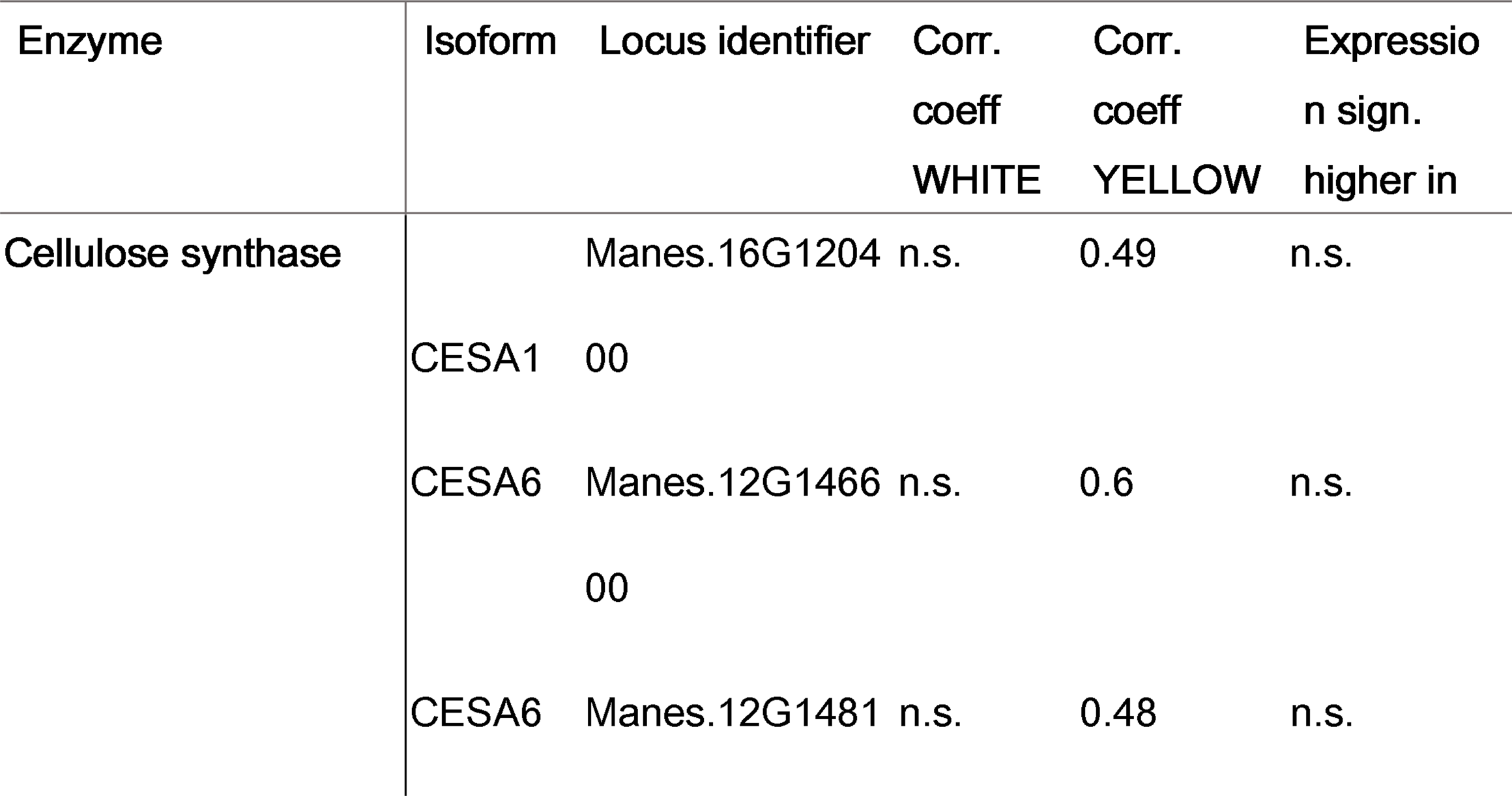
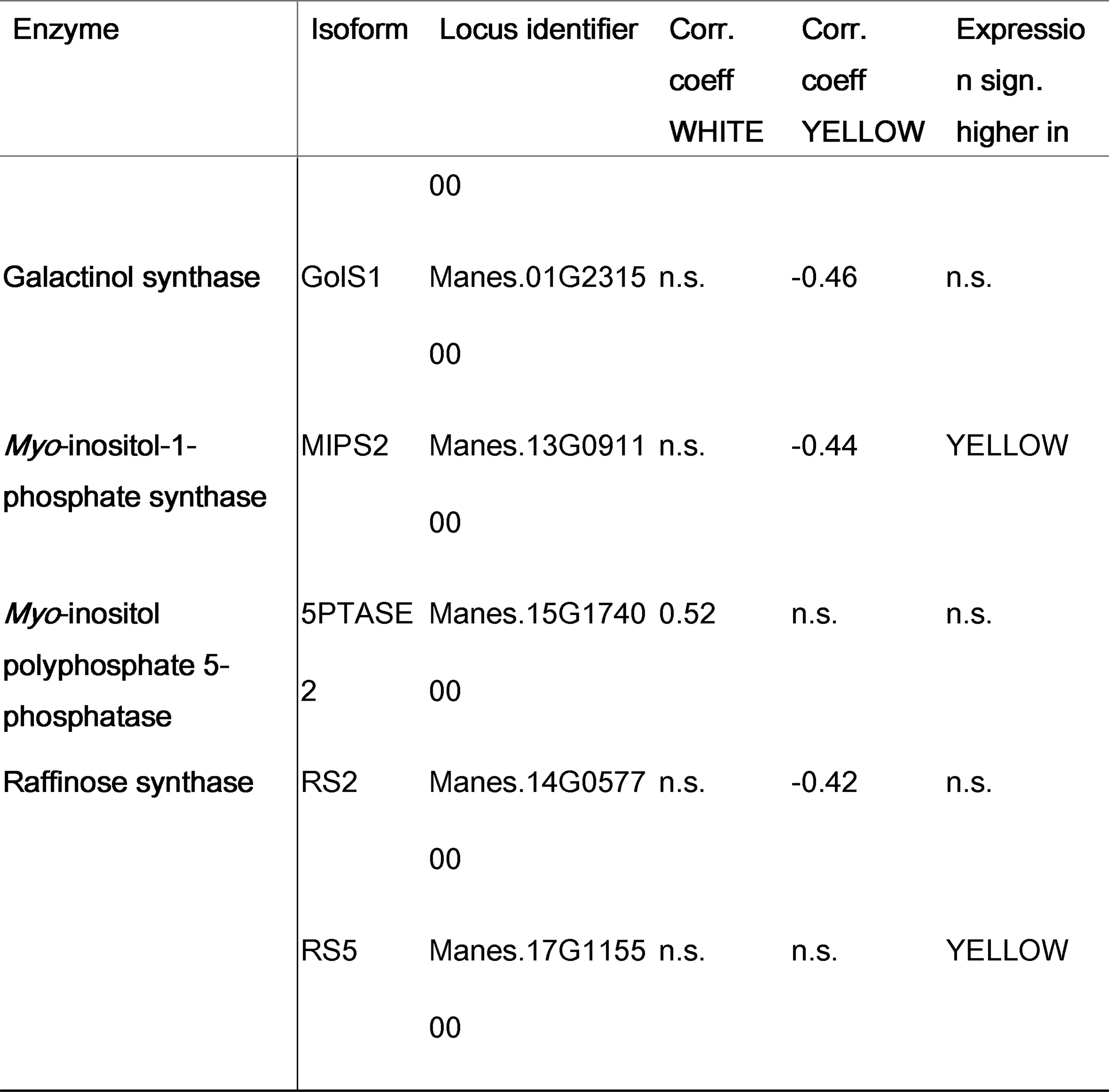
Enzymes related to the *myo*-inositol, raffinose and cell wall biosynthesis and identified as correlated to starch or significant differentially expressed between white and yellow genotypes.

| Enzyme | Isoform | Locus identifier | Corr.<br>coeff<br>WHITE | Corr.<br>coeff<br>YELLOW | Expressio<br>n sign.<br>higher in |
| --- | --- | --- | --- | --- | --- |
| Cellulose synthase |  | Manes.16G1204 | n.s. | 0.49 | n.s. |
|  | CESA1 | 00 |  |  |  |
|  | CESA6 | Manes.12G1466 | n.s. | 0.6 | n.s. |
|  |  | 00 |  |  |  |
|  | CESA6 | Manes.12G1481 | n.s. | 0.48 | n.s. |
|  |  | 00 |  |  |  |
| Galactinol synthase | GolS1 | Manes.01G2315 | n.s. | -0.46 | n.s. |
|  |  | 00 |  |  |  |
| *Myo*-inositol-1-phosphate synthase | MIPS2 | Manes.13G0911 | n.s. | -0.44 | YELLOW |
|  |  | 00 |  |  |  |
| *Myo*-inositol polyphosphate 5-phosphatase | 5PTASE2 | Manes.15G1740 | 0.52 | n.s. | n.s. |
|  |  | 00 |  |  |  |
| Raffinose synthase | RS2 | Manes.14G0577 | n.s. | -0.42 | n.s. |
|  |  | 00 |  |  |  |
|  | RS5 | Manes.17G1155 | n.s. | n.s. | YELLOW |
|  |  | 00 |  |  |  |

Endres & Tenhaken (2009) reported that *A. thaliana miox1 and miox2* knockout mutants, show a decreased, and overexpression of *MIOX4* an increased incorporation of MIOX-derived sugars into cell wall polymers. In addition, Wang *et al*. (2021) and Oleszkiewicz *et al*. (2021) hypothesized unknown signaling between cell wall and carotenoid biosynthesis. Wang *et al*. (2021) silenced *psy1* and *psy2* in *Nicotiana benthamiana* and *Nicotiana tabacum* L. varieties and observed not only a massive decrease in carotenoid biosynthesis, but also a negative effect on the expression of genes known to be involved in glucan, cellulose, pectin and galacturonan biosynthesis. Oleszkiewicz *et al*. (2021) generated among others, *psy2* mutants using CRISPR/Cas9 in carrot calli, and observed changes in the composition of arabinogalactan, pectin, and the extension in the cell walls of mutant plants.

To further investigate these observations, we asked the fundamental question if yellow cultivars possibly allocate a greater proportion of their carbon to cell wall components than their white counterparts. To address this question, we examined the relationship between starch-free debris and total dry yield in all genotypes studied. The results were both significant and revealing: the yellow genotypes had a significantly higher ratio of starch-free residue to total dry yield compared to the white varieties (t(48) = -2.3376, p=0.022; Figure 1F). This divergence not only underscores the differences in allocation strategies between the genotypes studied here, but might also provide a crucial insight which could support breeding decisions.

Oligosaccharides, such as raffinose and stachyose are formed from *myo*-inositol by the conversion from *myo*-inositol and UDP-D-Glucose to galactinol via GALACTINOL SYNTHASES (GolS). RAFFINOSE SYNTHASES (RS) then catalyze the transfer of the galactosyl unit from galactinol to sucrose, yielding raffinose. STACHYOSE SYNTHASES (STS) further transfers the galactosyl moiety from galactinol to galactose to produce stachyose (Loewus & Loewus, 1983; Loewus & Murthy, 2000). Yellow genotypes analyzed here do show higher abundance in all measured metabolites involved in *myo*-inositol and raffinose synthesis. As depicted in Figure 3 and 4, sucrose, *myo*-inositol and raffinose were detected with significantly higher abundance in yellow compared to white genotypes. The higher abundance of *myo-* inositol might additionally trigger raffinose and stachyose synthesis in the here analyzed yellow-fleshed cassava varieties. *RS1* was detected with a significantly higher expression in yellow compared to white genotypes (Figure 4, Table 5 and Supplementary material – Table S4-S6). Furthermore, *GolS1, RS1 and RS5* were identified with a significant negative correlation to starch in the yellow genotypes. Stachyose was not measured in the metabolite data analyzed here and in addition, *STACHYOSE SYNTHASE* (*STS*) could neither be detected to be differentially expressed between white and yellow genotypes nor to show a correlation to starch in either of the genotype groups (Figure 4, Table 5 and Supplementary material Table S4-S6).

**Figure 4:**
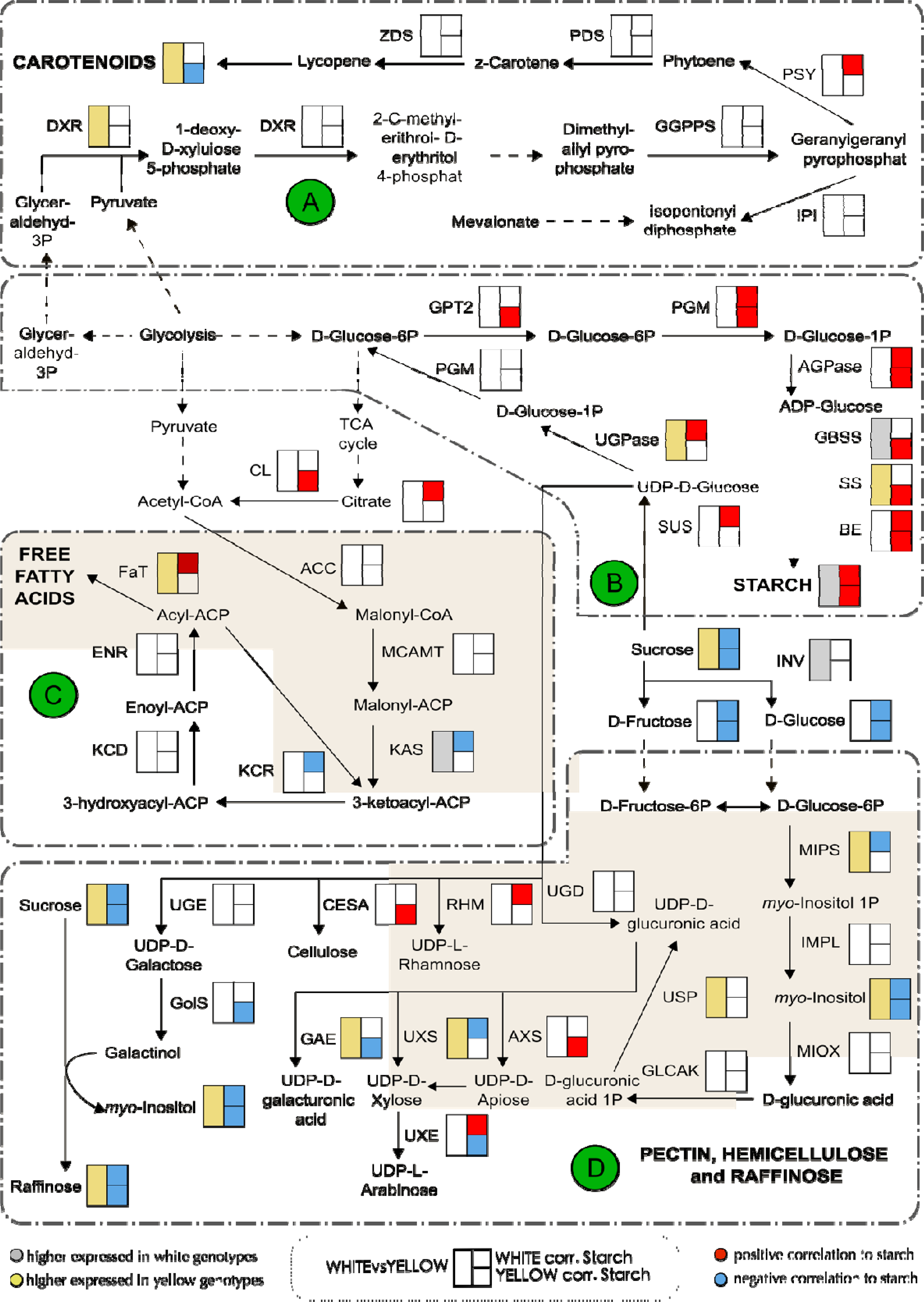
Gene expression patterns, correlation to starch and metabolite abundance. Model depicting the observed differences in gene (at least one for the corresponding group) or metabolite abundance between white and yellow genotypes and correlation to starch in either white or yellow genotypes for A) carotenoid biosynthesis B) sucrose breakdown and starch synthesis C) de novo fatty acid synthesis and D) synthesis of cell wall components and raffinose. Bigger left boxes shown besides the gene (*A. thaliana* homolog) or metabolite are filled gray if significantly higher abundant in white genotypes or yellow if significantly higher in yellow genotypes. Smaller boxes on the right show the significant correlation to starch with the upper box representing the correlation to starch in white genotypes and the lower for yellow genotypes. Red color indicates a positive correlation and blue a negative. Dashed lines refer to a non-direct synthesis path. Gene identifier, isoform, corresponding *A. thaliana* homolog, correlation coefficients, p-value and general abundance for all shown genes and metabolites are additionally provided in the Supplementary material. For detailed functional pathway description see main text.

Raffinose family oligosaccharides (RFOs) are known to be responsible for a wide variety of functions both in plants and the human diet. In recent years, RFOs have been heavily studied and identified to be involved in various stress responses in crops (Yan *et al*., 2022). For example, previous studies observed raffinose levels to be higher in plants showing increased cold-tolerance in *A. thaliana*, rice and sugar beet (Klotke *et al*., 2004; Morsy *et al*., 2007; Keller et al., 2021). Furthermore, RFOs are thought to be important for plant tolerance against heat stress, oxidative stress, drought, salt or biotic stresses (Yan *et al*., 2022). African *Manihot esculenta* varieties are subjected to multiple seasonal stressors like heat and drought during their growth period. Increased production of galactinol and raffinose through increased metabolic flux towards *myo*-inositol suggests that yellow-fleshed cassava genotypes might use it as a protective mechanism for the stress they are exposed to. However, independent of the stress source, and even though we could detect *GolS1*, *RS1* and *RS5* with a negative correlation to starch and a higher abundance in yellow genotypes, we can only hypothesize that the yellow genotypes might be better prepared for certain biotic or abiotic stresses.

Lastly, it is worth noting that for human consumption, some characteristics of RFOs need to be considered. Despite their reported potential health benefits, such as anti-allergic, anti-obesity, and anti-diabetic potentials (Kanwal *et al.,* 2023), RFOs are considered anti-nutritional factors. The human digestive system lacks the capacity to digest RFOs in the intestines, leading to their fermentation by gut microbiota. While this process develops intestinal symptoms like flatulence, abdominal cramps, diarrhea, and nausea, it is also important to recognize that these symptoms act as a barrier to RFO consumption in humans (Elango *et al*., 2022, Sanyal *et al*., 2023). Moreover, an increased proportion of RFOs in crops intended for human consumption may reduce the availability of metabolizable carbohydrates that serve as an energy source for humans. Consequently, the presence of RFOs in crops should be carefully managed to maintain an optimal balance between their benefits for crop cultivation and potential drawbacks for human consumers.

### Conclusion

Photosynthates that have been transported to the sink organs of plants can be metabolized into various carbon compounds like starch, fatty acids or cell wall components. The types and amounts of storage compounds differ significantly between plant species and genotypes and may heavily influence the value of the crop. Therefore, a key objective in many biotechnological approaches is to understand the mechanisms involved in carbon partitioning between metabolic pathways determining the final composition of the sink organ.

In this study, we investigated the differences in carbon partitioning between “white” and “yellow” cassava genotypes by analyzing a breeding panel with 49 African *Manihot esculenta* genotypes, with contrasting starch and carotenoid accumulation. With our experimental setup and analyses, we unified the three sources of data and were able to identify biochemical pathways that are linked to the crop’s phenotype, metabolome and transcriptome. In the genotypes under investigation, we examined (1) a negative correlation between starch and carotenoid content, (2) a metabolic separation between high starch and high carotenoid varieties, (3) a shift in gene expression from high starch to high carotenoid varieties. When taking the variety’s carotenoid levels into account, we observed (4) a positive correlation of genes involved in starch biosynthesis and starch content in both, white and yellow genotypes, (5) only minor differences in expression of genes involved in *de novo* fatty acid biosynthesis between white and yellow genotypes and almost no correlation to starch of those genes, (6) a significantly higher abundance and negative correlation to starch of almost all genes and all metabolites associated with *myo*-inositol, raffinose and the biosynthesis of building blocks for pectin and hemicellulose in yellow genotypes solely and (7) a significant higher starch-free residue to dry yield ratio in yellow-fleshed *Manihot esculenta* genotypes analyzed here. These results thus collectively enhance our understanding of the factors influencing the trade-off between starch, carotenoids and cell wall contents in cassava and will likely prove highly informative for future breeding strategies.

## Material and Methods

### Plant material, phenotyping and harvest

The population used in this study is composed of 49 African Cassava genotypes. The genotypes were planted as a preliminary yield trial in Ibadan (7°24’N, 3°54’E), Nigeria, from 2020 to 2021. Plots consisted of two rows with ten plants per plot, in a randomized complete block design with two replications. Spacings between rows and plants were 1 and 0.8m, respectively. Planting was performed from June to July 2020 and root harvesting was done the same time the following year. Phenotyping was done for all 49 genotypes for traits related to biotic stress, storage root quality (e.g., dry matter, starch and carotenoid content), and several other traits related to leaves, stems and roots.

### Sample preparation and RNA sequencing

For all 49 genotypes, storage root cross sections of six replicates per plot were harvested and pooled for further processing. All root samples were packed in aluminum foil, transferred to liquid nitrogen, and stored at -80°C until further processing.

RNA was extracted from all 49 samples (first replication) using the Spectrum plant Total RNA Kit. PolyA-selected library preparation and strand unspecific paired-end sequencing was provided as a custom service (Illumina NovaSeq 6000; Novogene, Cambridge, UK).

### Metabolite measurements

The same material as used for RNA extraction was used for metabolome analysis using gas chromatography-mass spectrometry (GC-MS) analysis and was processed as described in Rosado-Souza et al. (2019). Data was reported according to recently updated reporting standards for metabolomics (Alseekh et al., 2021; Supplementary material Table S2).

### Data pre-processing

Since the linear relationship between the intensity of yellow color in cassava storage roots and the carotenoid content (Pearson’s r ∼0.8) is already well studied (Iglesias *et al*., 1997; Chávez *et al*., 2005; Akinwale *et al*., 2010; Sánchez *et al*., 2014), we used yellowness as a proxy for carotenoid content. To avoid subjectivity by measuring the yellowness visually, a chromameter was used to assess lightness (L*), red and green (a*, positive = red, negative = green) as well as yellow and blue (b*, positive = yellow, negative = blue) color values using the Commission Internationale de l’Éclairage (CIELAB) method. Genotypes with a positive Chromameter b* ≤ 25 were identified as low carotenoid content genotypes (referred to as white genotypes), whereas genotypes showing a b* > 25 were classified as high carotenoid content genotypes (referred to as yellow genotypes). In the whole manuscript, carotenoid content refers to the chromameter b* values as a measure for carotenoid content. For starch content analyses, the percentage of starch per genotype analyzed with Near infrared (NIR) spectroscopy was used.

### Bioinformatic analyses

Raw sequence reads were inspected using FastQC v.0.11.9 (Andrews, 2010) and MultiQC v.1.13 (Ewels *et al*., 2016). Using BBMap v.38.97 (Bushnell, 2014) the reads were filtered for N-reads, common contaminants, and known adaptor sequences, subsequently. Reads were additionally filtered based on read quality (≥30) and length (≥35). Quality reports for checking trimming success were generated using FastQC and MultiQC. Adapter- and quality-trimmed RNAseq reads were mapped to the *Manihot esculenta* v8.1 reference genome, using STAR (v.2.7.10a) (Dobin *et al*., 2013). *Manihot esculenta* v8.1 reference genome and corresponding annotations were downloaded from Phytozome (available under: https://phytozome-next.jgi.doe.gov/info/Mesculenta_v8_1) and served as reference genome during any analyses (Goodstein *et al*., 2012). Uniquely mapped transcriptomic reads were quantified on gene-level using featureCounts v.2.0.3 (Liao *et al*., 2014).

For sequence analysis, a variant calling, including all genotypes, was performed using the BCFtools v.1.15.1 *call* function (Li, 2011). Subsequent filters were applied using VCFtools v.0.1.17 (Danecek *et al*., 2011). Filtering parameters were adjusted according to minimum quality (20), depth (≥60), minor allele frequency (0.05), maximum number of genotypes allowed with missing value for called position (10% percent), and biallelic single nucleotide polymorphisms (SNPs).

Differential expression analysis between white and yellow varieties was conducted using DESeq2 (Love *et al*., 2014) in R. DEGs were subsequently filtered for average gene-level quantified read depth of ≥10. Average read depth was calculated for white and yellow genotypes separately to avoid falsely excluding genes which are solely expressed in either white or yellow genotypes. P-value adjustment was performed using the false discovery rate (FDR), accepting genes as significant differentially expressed with a threshold of FDR ≤.01.

Depth filtered (≥10), variance-stabilizing transformed high quality reads were used to identify genes which are significantly correlated to starch, while controlling for carotenoid content as a confounding effect, in the white and the yellow genotypes separately by performing a partial correlation analysis using the ppcor package (Kim, 2015) in R. Partial correlation, quantifies the degree of association between two variables (here gene expression and starch content), while adjusting for potential confounding effects of a third variable (carotenoid content). In simpler terms, this can be understood as examining the relationship between gene expression and starch while holding carotenoid levels constant. Transcripts showing p ≤.05 and r ≥ |0.3| were considered being significantly correlated. If transcriptional correlation to starch is mentioned in the manuscript, we are always referring to a correlation to starch including carotenoid content as a confounding variable. Subsequently, significant correlated genes were used to perform a functional enrichment analysis using the one-sided Fisher’s exact test provided in the clusterProfiler package (Yu *et al*., 2012) in R. Genes were functionally categorized using KEGG orthology (KO) terms. *Manihot esculenta* genes were described by their best *A. thaliana* hit based on BLASTP similarity. Pathway construction and identification of gene interactions were constructed through publications and publicly available literature and are listed in the corresponding sections.

In order to test whether yellow cultivars allocate a different proportion of their carbon resources to cell wall components than their white counterparts, we calculated the ratio of starch-free debris to total dry yield content as followed:

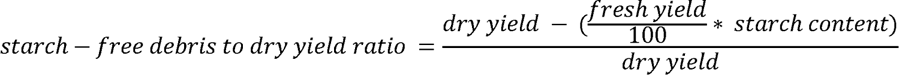

where dry yield, fresh yield and starch content represent the phenotypic observations for the corresponding variety. For subsequent statistical significance testing, the ratio of all 49 samples of both replications (different plot locations on the field) were collapsed to single best linear unbiased estimator (BLUE) for white and yellow varieties. BLUE was calculated using the lmer function of the lme4 package in R (Bates *et al*., 2014) as follows:

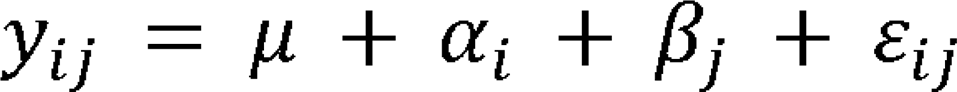

where *y*_*ij*_ represents the ratio observed for variety *i* at plot location *j*, *µ* is the grad mean, *a*_*i*_ is the random effect for variety, *{J*_*j*_ representing the fixed effects term for plot location and *E*_*ij*_ the residual variance. For investigations including the metabolites, a z-score transformation of log values of the measured metabolites was performed and used for all subsequent analyses. Metabolites showing p ≤.05 and r ≥ |0.3| were accepted as being significantly correlated to starch.

All analyses outside of R were performed under Linux (Ubuntu v20.04.4). Analyses in R were performed under macOS (Catalina v10.15.7) and R version 4.1.0. For a detailed listing of all packages and corresponding versions used in R, see Supplementary material - R Session info. For visualization, if not otherwise specified, plots were generated using ggplot (Wickham, 2016) in R and finalized using Inkscape v1.0.2 (*Inkscape*, 2020).

## Data availability

All used commands and scripts are available on github (https://github.com/Division-of-Biochemistry-Publications/CASS).

Raw sequencing reads have been deposited to NCBI’s Sequence Read Archive (submission ID: SUB13965463) under BioProject ID PRJNA1039569, available at https://www.ncbi.nlm.nih.gov/sra/PRJNA1039569.

## Acknowledgments

We thank Otilia Ciobotea and Michaela Reiser for their excellent technical assistance.

## Author contributions

WZ, US, IYR designed the research. IYR and AMD designed and executed the field trials. AS processed the sample material. ARF and LRS conceived and performed the metabolic analyses. HEN and BP conceived and performed the sugar measurements. SG, DR, SR performed the analysis. WZ and US organized and supervised the work. SG wrote the manuscript with contributions of all authors. All authors have read and approved the submitted version of the manuscript.

## Legends for supporting information

**Table S1:** All measured metabolites and respective assignment as white or yellow variety.

**Table S2:** Metabolite reporting checklist.

**Table S3:** Genotype distribution for all genotypes for *PSY2* SNP position 31960252.

**Table S4:** List of all significant differentially expressed genes between white and yellow genotypes.

**Table S5:** Listing of all genes significantly correlated to starch (including carotenoid content as confounding effect) in one of the two genotype groups.

**Table S6:** Listing of all genes significant differentially expressed between white and yellow genotypes, significantly correlated to starch in one of the genotype groups, or specifically mentioned in the main text of the manuscript.

**Figure S1:**
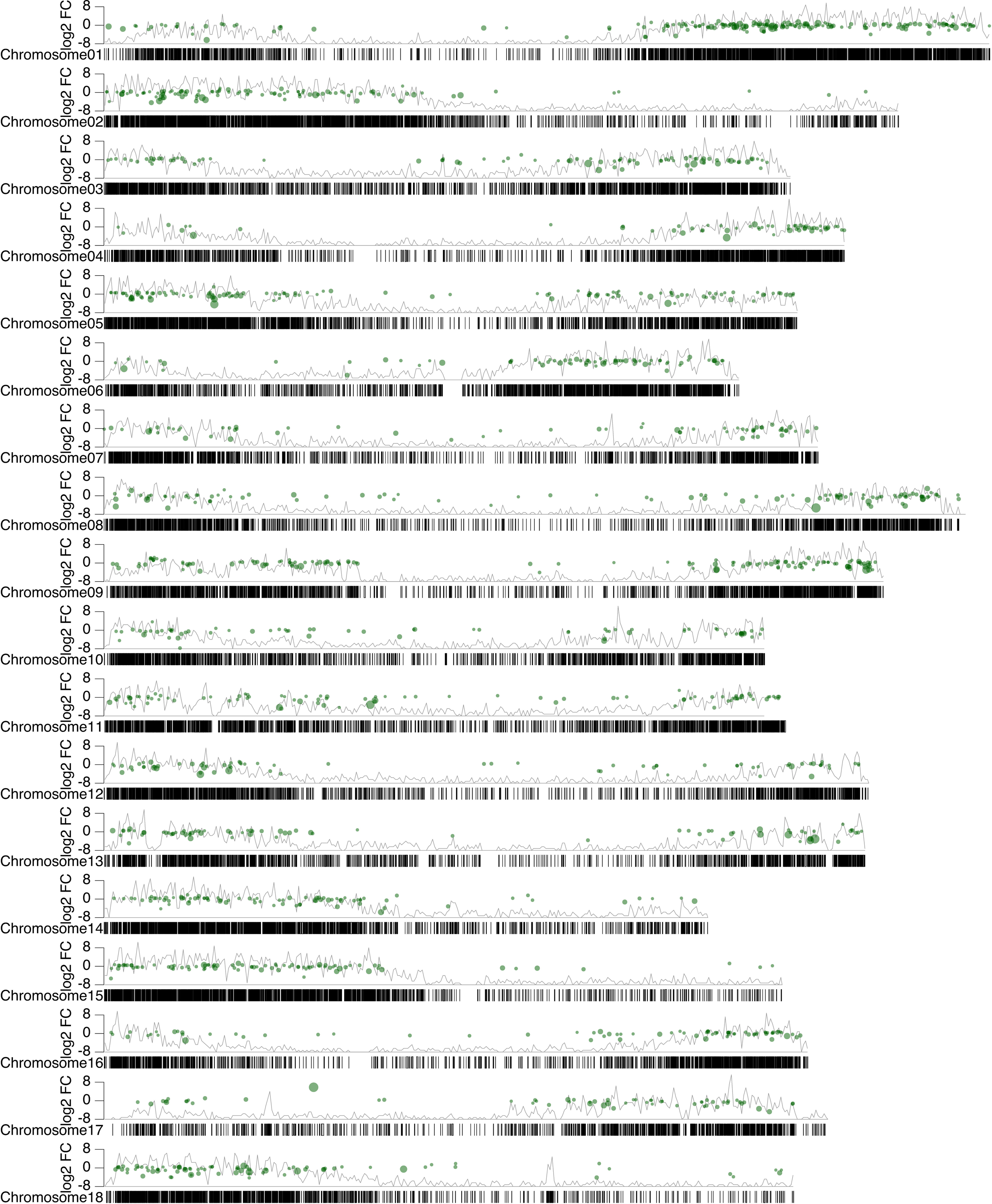
Chromosomal distribution of all detected significantly differential expressed genes between white and yellow genotypes across all 18 *Manihot esculenta* chromosomes.

**Figure S2:**
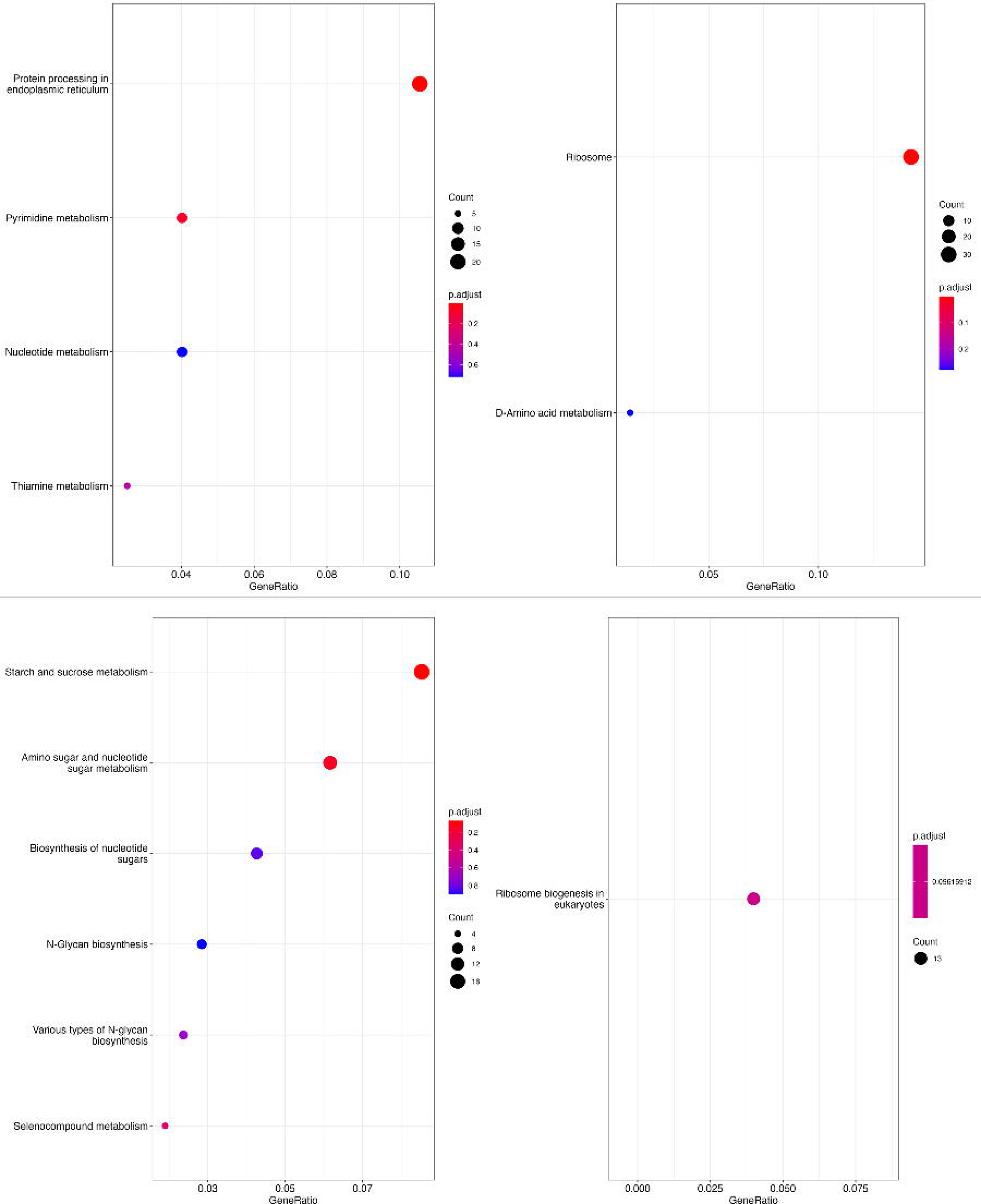
Functional enrichment analysis for genes showing significant correlation to starch after correcting for carotenoid content as confounding effect.

## References

A. Guschina, I., & Harwood, J. L. (2007). Complex lipid biosynthesis and its manipulation in plants. In P. Ranalli (Ed.), Improvement of Crop Plants for Industrial End Uses (pp. 253–279). Springer Netherlands.

Akinwale, M. G., Aladesanwa, R. D., Akinyele, B. O., Dixon, A. G. O., & Odiyi, A. C. (2010). Inheritance of ß-carotene in cassava (Manihot esculenta crantza). *Vol.* 2(10), 198–201.

Alseekh, S., Aharoni, A., Brotman, Y., Contrepois, K., D’Auria, J., Ewald, J., Ewald, J. C., Fraser, P. D., Giavalisco, P., Hall, R. D., Heinemann, M., Link, H., Luo, J., Neumann, S., Nielsen, J., Perez de Souza, L., Saito, K., Sauer, U., Schroeder, F. C., Schuster, S., Siuzdak, G., Skirycz, A., Sumner, L. W., Snyder, M. P., Tang, H., Tohge, T., Wang, Y., Wen, W., Wu, S., Xu, G., Zamboni, N. & Fernie, A. R. (2021). Mass spectrometry-based metabolomics: a guide for annotation, quantification and best reporting practices. Nature methods, 18(7), 747–756.

Andrews, S. R. (2010). FastQC Available at: http://www.bioinformatics.babraham.ac.uk/projects/fastqc/

Awotide, B., Abdoulaye, T., Alene, A., & Manyong, V. M. (2014). Assessing the extent and determinants of adoption of improved cassava varieties in south-western Nigeria. Journal of Development and Agricultural Economics, 6(9), 376–385.

Bates, D., Mächler, M., Bolker, B., & Walker, S. (2014). Fitting Linear Mixed-Effects Models using lme4.

Beyene, G., Solomon, F. R., Chauhan, R. D., Gaitán-Solis, E., Narayanan, N., Gehan, J., Siritunga, D., Stevens, R. L., Jifon, J., Van Eck, J., Linsler, E., Gehan, M., Ilyas, M., Fregene, M., Sayre, R. T., Anderson, P., Taylor, N. J., & Cahoon, E. B. (2018). Provitamin A biofortification of cassava enhances shelf life but reduces dry matter content of storage roots due to altered carbon partitioning into starch. Plant Biotechnology Journal, 16(6), 1186–1200.

Bushnell, B. (2014). BBMap short read aligner, and other bioinformatic tools. In BBMap short read aligner, and other bioinformatic tools. Available at: https://sourceforge.net/projects/bbmap/

Cai, J., Xue, J., Zhu, W., Luo, X., Lu, X., Xue, M., Wie, Z., Cai, Y., Ou, W., Li, K., An, F. & Chen, S. (2023). Integrated Metabolomic and Transcriptomic Analyses Reveals Sugar Transport and Starch Accumulation in Two Specific Germplasms of Manihot esculenta Crantz. International Journal of Molecular Sciences, 24(8), 7236.

Carvalho, L. J., Filho, J. F., Anderson, J. V., Figueiredo, P. W., & Chen, S. (2018). Storage Root of Cassava: Morphological Types, Anatomy, Formation, Growth, Development and Harvest Time. In V. Waisundara (Ed.), Cassava. InTech.

Chavez, A. L., Bedoya, J. M., Sánchez, T., Iglesias, C., Ceballos, H., & Roca, W. (2000). Iron, carotene, and ascorbic acid in cassava roots and leaves. Food and Nutrition Bulletin, 21(4), 410–413.

Chávez, A. L., Sánchez, T., Jaramillo, G., Bedoya, J. M., Echeverry, J., Bolaños, E. A., Ceballos, H., & Iglesias, C. A. (2005). Variation of quality traits in cassava roots evaluated in landraces and improved clones. Euphytica, 143(1–2), 125– 133.

Cunningham, F. X., & Gantt, E. (1998). GENES AND ENZYMES OF CAROTENOID BIOSYNTHESIS IN PLANTS. Annual Review of Plant Physiology and Plant Molecular Biology, 49(1), 557–583.

Danecek, P., Auton, A., Abecasis, G., Albers, C. A., Banks, E., DePristo, M. A., Handsaker, R. E., Lunter, G., Marth, G. T., Sherry, S. T., McVean, G., Durbin, R., & 1000 Genomes Project Analysis Group. (2011). The variant call format and VCFtools. Bioinformatics, 27(15), 2156–2158.

Dobin, A., Davis, C. A., Schlesinger, F., Drenkow, J., Zaleski, C., Jha, S., Batut, P., Chaisson, M., & Gingeras, T. R. (2013). STAR: Ultrafast universal RNA-seq aligner. Bioinformatics, 29(1), 15–21.

Elango, D., Rajendran, K., Van der Laan, L., Sebastiar, S., Raigne, J., Thaiparambil, N. A., Haddad, N. E., Raja, B., Wang, W., Ferela, A., Chiteri, K. O., Thudi, M., Varshney, R. K., Chopra, S., Singh A. & Singh, A. K. (2022). Raffinose family oligosaccharides: friend or foe for human and plant health?. Frontiers in Plant Science, 13, 829118.

Endres, S., & Tenhaken, R. (2009). Myoinositol Oxygenase Controls the Level of Myoinositol in Arabidopsis, But Does Not Increase Ascorbic Acid. Plant Physiology, 149(2), 1042–1049.

Esuma, W., Herselman, L., Labuschagne, M. T., Ramu, P., Lu, F., Baguma, Y., Buckler, E. S., & Kawuki, R. S. (2016). Genome-wide association mapping of provitamin A carotenoid content in cassava. Euphytica, 212(1), 97–110.

Ewels, P., Magnusson, M., Lundin, S., & Käller, M. (2016). MultiQC: Summarize analysis results for multiple tools and samples in a single report. Bioinformatics, 32(19), 3047–3048.

FAO. (2023). Crops and livestock products. Available at: https://www.fao.org/faostat/en/#data/QCL

Farré, G., Sanahuja, G., Naqvi, S., Bai, C., Capell, T., Zhu, C., & Christou, P. (2010). Travel advice on the road to carotenoids in plants. Plant Science, 179(1–2), 28–48.

Figueroa, C. M., Asencion Diez, M. D., Ballicora, M. A., & Iglesias, A. A. (2022). Structure, function, and evolution of plant ADP-glucose pyrophosphorylase. Plant Molecular Biology, 108(4–5), 307–323.

Goodstein, D. M., Shu, S., Howson, R., Neupane, R., Hayes, R. D., Fazo, J., Mitros, T., Dirks, W., Hellsten, U., Putnam, N. & Rokhsar, D. S. (2012). Phytozome: a comparative platform for green plant genomics. Nucleic acids research, 40(D1), D1178–D1186.

Harwood, J. L. (1996). Recent advances in the biosynthesis of plant fatty acids. Biochimica et Biophysica Acta (BBA) - Lipids and Lipid Metabolism, 1301(1– 2), 7–56.

Hefferon, K. (2015). Nutritionally Enhanced Food Crops; Progress and Perspectives. International Journal of Molecular Sciences, 16(2), 3895–3914.

Iglesias, C., Mayer, J., Chavez, L., & Calle, F. (1997). Genetic potential and stability of carotene content in cassava roots. Euphytica, 94(3), 367–373.

Inkscape. (2020). Available at: https://inkscape.org/release/inkscape-1.2.2/

Jonik, C., Sonnewald, U., Hajirezaei, M. R., Flügge, U. I., & Ludewig, F. (2012). Simultaneous boosting of source and sink capacities doubles tuber starch yield of potato plants. Plant Biotechnology Journal, 10(9), 1088–1098.

Ke, J., Behal, R. H., Back, S. L., Nikolau, B. J., Wurtele, E. S., & Oliver, D. J. (2000). The Role of Pyruvate Dehydrogenase and Acetyl-Coenzyme A Synthetase in Fatty Acid Synthesis in Developing Arabidopsis Seeds. Plant Physiology, 123(2), 497–508.

Keller, I., Müdsam, C., Rodrigues, C. M., Kischka, D., Zierer, W., Sonnewald, U., Harms, K., Czarnecki, O., Fiedler-Wiechers, K., Koch, W., Neuhaus, H. E., Ludewig, F., & Pommerrenig, B. (2021). Cold-Triggered Induction of ROS-and Raffinose Metabolism in Freezing-Sensitive Taproot Tissue of Sugar Beet. Frontiers in Plant Science, 12, 715767.

Kim, S. (2015). ppcor: An R Package for a Fast Calculation to Semi-partial Correlation Coefficients. Communications for Statistical Applications and Methods, 22(6), 665–674.

Klaus, D., Ohlrogge John, B., Neuhaus, H. E., & Dörmann, P. (2004). Increased fatty acid production in potato by engineering of acetyl-CoA carboxylase. Planta, 219(3).

Klotke, J., Kopka, J., Gatzke, N., & Heyer, A. G. (2004). Impact of soluble sugar concentrations on the acquisition of freezing tolerance in accessions of Arabidopsis thaliana with contrasting cold adaptation - evidence for a role of raffinose in cold acclimation: Cold acclimation of Arabidopsis accessions. Plant, Cell & Environment, 27(11), 1395–1404.

Lamm, C. E., Rabbi, I. Y., Medeiros, D. B., Rosado-Souza, L., Pommerrenig, B., Dahmani, I., Rüscher, D., Hofmann, J., Van Doorn, A. M., Schlereth, A., Neuhaus, H. E., Fernie, A. R., Sonnewald, U., & Zierer, W. (2023). Efficient sugar utilization and transition from oxidative to substrate-level phosphorylation in high starch storage roots of African cassava genotypes. The Plant Journal, tpj.16357.

Li, H. (2011). A statistical framework for SNP calling, mutation discovery, association mapping and population genetical parameter estimation from sequencing data. Bioinformatics, 27(21), 2987–2993.

Liao, Y., Smyth, G. K., & Shi, W. (2014). featureCounts: An efficient general purpose program for assigning sequence reads to genomic features. Bioinformatics, 30(7), 923–930.

Loewus, F. A., & Loewus, M. W. (1983). myo -Inositol: Its Biosynthesis and Metabolism. Annual Review of Plant Physiology, 34(1), 137–161.

Loewus, F. A., & Murthy, P. P. N. (2000). Myo-Inositol metabolism in plants. Plant Science, 150(1), 1–19.

Love, M. I., Huber, W., & Anders, S. (2014). Moderated estimation of fold change and dispersion for RNA-seq data with DESeq2. Genome Biology, 15(12).

Luo, X., An, F., Xue, J., Zhu, W., Wei, Z., Ou, W., Li, K., Chen, S. & Cai, J. (2023). Integrative analysis of metabolome and transcriptome reveals the mechanism of color formation in cassava (Manihot esculenta Crantz) leaves. Frontiers in Plant Science, 14, 1181257.

Mehdi, R., Lamm, C. E., Bodampalli Anjanappa, R., Müdsam, C., Saeed, M., Klima, J., Kraner, M. E., Ludewig, F., Knoblauch, M., Gruissem, W., Sonnewald, U., & Zierer, W. (2019). Symplasmic phloem unloading and radial post-phloem transport via vascular rays in tuberous roots of Manihot esculenta. Journal of Experimental Botany, 70(20), 5559–5573.

Morsy, M. R., Jouve, L., Hausman, J.-F., Hoffmann, L., & Stewart, J. McD. (2007). Alteration of oxidative and carbohydrate metabolism under abiotic stress in two rice (Oryza sativa L.) genotypes contrasting in chilling tolerance. Journal of Plant Physiology, 164(2), 157–167.

Munyikwa, T. R. I., Langeveld, S., Salehuzzaman, S. N. I. M., Jacobsen, E., & Visser, R. G. F. (1997). Cassava starch biosynthesis: New avenues for modifying starch quantity and quality. Euphytica, 96(1), 65–75.

Nassar, N., & Ortiz, R. (2010). Breeding Cassava to Feed the Poor. Scientific American, 302(5), 78–84.

Ogbonna, A. C., Ramu, P., Esuma, W., Nandudu, L., Morales, N., Powell, A., Kawuki, R., Bauchet, G., Jannink, J. & Mueller, L. A. (2021). A population based expression atlas provides insights into disease resistance and other physiological traits in cassava (Manihot esculenta Crantz). Scientific Reports, 11(1), 23520.

Olayide, P., Alexandersson, E., Tzfadia, O., Lenman, M., Gisel, A., & Stavolone, L. (2023). Transcriptome and metabolome profiling identify factors potentially involved in pro-vitamin A accumulation in cassava landraces. Plant Physiology and Biochemistry, 199, 107713.

Oleszkiewicz, T., Klimek-Chodacka, M., Kruczek, M., Godel-Jędrychowska, K., Sala, K., Milewska-Hendel, A., Zubko, M., Kurczyńska, E., Qi, Y., & Baranski, R. (2021). Inhibition of Carotenoid Biosynthesis by CRISPR/Cas9 Triggers Cell Wall Remodelling in Carrot. International Journal of Molecular Sciences, 22(12), 6516.

Rabbi, I. Y., Udoh, L. I., Wolfe, M., Parkes, E. Y., Gedil, M. A., Dixon, A., Ramu, P., Jannink, J., & Kulakow, P. (2017). Genome-Wide Association Mapping of Correlated Traits in Cassava: Dry Matter and Total Carotenoid Content. The Plant Genome, 10(3).

Rabbi, I. Y., Kayondo, S. I., Bauchet, G., Yusuf, M., Aghogho, C. I., Ogunpaimo, K., Uwugiaren, R., Smith, I. A., Peteti, P., Agbona, A., Parkes, E., Lydia, E., Wolfe, M., Jannink, J., Egesi, C. & Kulakow, P. (2022). Genome-wide association analysis reveals new insights into the genetic architecture of defensive, agro-morphological and quality-related traits in cassava. Plant Molecular Biology, 109(3), 195–213.

Rawsthorne, S. (2002). Carbon flux and fatty acid synthesis in plants. Progress in Lipid Research, 41(2), 182–196.

Rosado-Souza, L., David, L. C., Drapal, M., Fraser, P. D., Hofmann, J., Klemens, P. A. W., Ludewig, F., Neuhaus, H. E., Obata, T., Perez-Fons, L., Schlereth, A., Sonnewald, U., Stitt, M., Zeeman, S. C., Zierer, W., & Fernie, A. R. (2019). Cassava Metabolomics and Starch Quality. Current Protocols in Plant Biology, 4(4).

Sánchez, T., Ceballos, H., Dufour, D., Ortiz, D., Morante, N., Calle, F., Zum Felde, T., Domínguez, M., & Davrieux, F. (2014). Prediction of carotenoids, cyanide and dry matter contents in fresh cassava root using NIRS and Hunter color techniques. Food Chemistry, 151, 444–451.

Sanyal, R., Kumar, S., Pattanayak, A., Kar, A., & Bishi, S. K. (2023). Optimizing raffinose family oligosaccharides content in plants: A tightrope walk. Frontiers in Plant Science, 14, 1134754.

Talsma, E. F., Borgonjen-van Den Berg, K. J., Melse-Boonstra, A., Mayer, E. V., Verhoef, H., Demir, A. Y., Ferguson, E. L., Kok, F. J., & Brouwer, I. D. (2018). The potential contribution of yellow cassava to dietary nutrient adequacy of primary-school children in Eastern Kenya; the use of linear programming. Public Health Nutrition, 21(2), 365–376.

Telengech, P. K., Maling’a, J. N., Nyende, A. B., Gichuki, S. T., & Wanjala, B. W. (2015). Gene expression of beta carotene genes in transgenic biofortified cassava. 3 Biotech, 5(4), 465–472.

Walter, M. H., & Strack, D. (2011). Carotenoids and their cleavage products: Biosynthesis and functions. Natural Product Reports, 28(4), 663.

Wang, Z., Zhang, L., Dong, C., Guo, J., Jin, L., Wei, P., Li, F., Zhang, X., & Wang, R. (2021). Characterization and functional analysis of phytoene synthase gene family in tobacco. BMC Plant Biology, 21(1), 32.

Welsch, R., Arango, J., Bär, C., Salazar, B., Al-Babili, S., Beltrán, J., Chavarriaga, P., Ceballos, H., Tohme, J., & Beyer, P. (2010). Provitamin A Accumulation in Cassava (Manihot esculenta) Roots Driven by a Single Nucleotide Polymorphism in a Phytoene Synthase Gene. The Plant Cell, 22(10), 3348– 3356.

Wickham, H. (2016). ggplot2: Elegant Graphics for Data Analysis (2nd ed. 2016). Springer International Publishing□: Imprint: Springer.

Wilson, M. C., Mutka, A. M., Hummel, A. W., Berry, J., Chauhan, R. D., Vijayaraghavan, A., Taylor, N. J., Voytas, D. F., Chitwood, D. H. & Bart, R. S. (2017). Gene expression atlas for the food security crop cassava. New Phytologist, 213(4), 1632–1641.

Wright, L. P., Rohwer, J. M., Ghirardo, A., Hammerbacher, A., Ortiz-Alcaide, M., Raguschke, B., Schnitzler, H. P., Gershenzon, J. & Phillips, M. A. (2014). Deoxyxylulose 5-phosphate synthase controls flux through the methylerythritol 4-phosphate pathway in Arabidopsis. Plant Physiology, 165(4), 1488–1504.

Xiao, L., Cao, S., Shang, X., Xie, X., Zeng, W., Lu, L., Kong, Q. & Yan, H. (2021). Metabolomic and transcriptomic profiling reveals distinct nutritional properties of cassavas with different flesh colors. Food Chemistry: Molecular Sciences, 2, 100016.

Yan, S., Liu, Q., Li, W., Yan, J., & Fernie, A. R. (2022). Raffinose Family Oligosaccharides: Crucial Regulators of Plant Development and Stress Responses., 16(5), 284–287.

Zeeman, S. C., Kossmann, J., & Smith, A. M. (2010). Starch: Its Metabolism, Evolution, and Biotechnological Modification in Plants. Annual Review of Plant Biology, 61(1), 209–234.

Zhang, L., Häusler, R. E., Greiten, C., Hajirezaei, M. R., Haferkamp, I., Neuhaus, H. E., Flügge, U. I. & Ludewig, F. (2008). Overriding the co-limiting import of carbon and energy into tuber amyloplasts increases the starch content and yield of transgenic potato plants. Plant biotechnology journal, 6(5), 453–464.

Zhu, F. (2015). Composition, structure, physicochemical properties, and modifications of cassava starch. Carbohydrate Polymers, 122, 456–480.

