## Supplementarial material for "Carbon usage in yellow-fleshed *Manihot esculenta* storage roots shifts from starch biosynthesis to cell wall and raffinose biosynthesis via the *myo*-inositol pathway"

**All figures and tables that can be meaningfully presented in this document are shown below. Tables that are too large to be displayed in this document are described in the following and titled and attached in separate files with their corresponding name.**

**Supplementary material - Table S1 and S2**

All measured metabolites for all genotypes under investigation including the assignment to the respective genotype group - see file “Measured_metabolites.txt”. Metabolite reporting standards see file “Metabolite_reporting_checklist.xlsx”.

**Supplementary material - Table S3**

Genotype distribution for all genotypes for *PSY2* SNP position 31960252. All white genotypes show homozygosity (0/0) for the reference allele (C), whereas all yellow genotypes show either heterozygosity (0/1) for the corresponding location or homozygosity (1/1) for the alternative allele (A).

| **genotype for *PSY2*** | **detected b*** | **categorized as** |
| --- | --- | --- |
| 0/0 | 14.99 | WHITE |
| 0/0 | 16.355 | WHITE |
| 0/0 | 17.02 | WHITE |
| 0/0 | 17.17 | WHITE |
| 0/0 | 17.35 | WHITE |
| 0/0 | 17.79 | WHITE |
| 0/0 | 17.8 | WHITE |
| 0/0 | 17.875 | WHITE |
| 0/0 | 17.985 | WHITE |
| 0/0 | 18.085 | WHITE |
| 0/0 | 18.22 | WHITE |
| 0/0 | 18.285 | WHITE |
| 0/0 | 18.43 | WHITE |
| 0/0 | 18.985 | WHITE |
| 0/0 | 19.18 | WHITE |
| 0/0 | 19.575 | WHITE |
| 0/0 | 19.66 | WHITE |
| 0/0 | 20.315 | WHITE |
| 0/0 | 20.37 | WHITE |
| 0/0 | 22.275 | WHITE |
| 0/0 | 22.365 | WHITE |
| 0/0 | 22.485 | WHITE |
| 0/1 | 27.98 | YELLOW |
| 0/1 | 29.075 | YELLOW |
| 0/1 | 30.615 | YELLOW |
| 0/1 | 32.26 | YELLOW |
| 0/1 | 33.575 | YELLOW |
| 0/1 | 34.555 | YELLOW |
| 0/1 | 36.67 | YELLOW |
| 0/1 | 37.055 | YELLOW |
| 0/1 | 38.385 | YELLOW |
| 1/1 | 33.185 | YELLOW |
| 1/1 | 34.26 | YELLOW |
| 1/1 | 38.83 | YELLOW |
| 1/1 | 40.465 | YELLOW |
| 1/1 | 40.525 | YELLOW |
| 1/1 | 40.565 | YELLOW |
| 1/1 | 40.605 | YELLOW |
| 1/1 | 41.2 | YELLOW |
| 1/1 | 42.035 | YELLOW |
| 1/1 | 42.235 | YELLOW |
| 1/1 | 43.1 | YELLOW |
| 1/1 | 43.685 | YELLOW |
| 1/1 | 43.87 | YELLOW |
| 1/1 | 44.29 | YELLOW |
| 1/1 | 48.06 | YELLOW |
| 1/1 | 48.7 | YELLOW |
| 1/1 | 48.875 | YELLOW |

**Supplementary material - Table S4**

Differential expression analysis results between white and yellow genotypes. See file “DE_WHITEvsYELLOW.txt”. A positive log2FoldChange and an adjusted p-value <= .05 represent genes with a significantly higher abundance in white genotypes, whereas a positive log2FoldChange and an adjusted p-value <= .05 represents all genes with a higher abundance in yellow genotypes. All Gene IDs were functionally annotated, therefore some gene IDs might be doubled because one gene ID in *Manihot esculenta* can refer to more than one *A. thaliana* locus ID and vice versa. For the functional annotation method see main manuscript.

**Supplementary material - Table S5**

Listing of all genes significantly correlated to starch (including carotenoid content as covariate) in one of the two genotype groups. See file “Genes_correlated_to_starch.xlsx”. Sheet “WHITE” is representing genes showing a significant correlation to starch in the white genotypes. “YELLOW” is representing genes showing a significant correlation to starch in the yellow genotypes. All Gene IDs were functionally annotated, therefore some gene IDs might be doubled because one gene ID in *Manihot esculenta* can refer to more than one *A. thaliana* locus ID and vice versa. For the functional annotation method see main manuscript.

**Supplementary material - Table S6**

Listing of all genes significant differentially expressed between white and yellow genotypes, significantly correlated to starch in one of the genotype groups, or specifically mentioned in the main text of the manuscript. See file “GOIs.txt”. The first column is depicting the *Manihot esculenta* locus identifier. The next four columns refer to the correlation coefficient and p-value in either white or yellow genotypes. Column F and G representing the adjusted p-value and the log2FC for that locus comparing white and yellow genotypes. “functional annotation”, “abbreviation” and “*A. thaliana* homolog” showing the corresponding functional annotation based on the best *A. thaliana* BLASTP hit. The last two columns “expressed in WHITE” and “expressed in YELLOW” represent, if the locus was detected as expressed (normalized read counts > 10). All Gene IDs were functionally annotated, therefore some gene IDs might be doubled because one gene ID in *Manihot esculenta* can refer to more than one *A. thaliana* locus ID and vice versa. For functional annotation and expression detection method see main manuscript.

**Supplementary material - Figure S1**

Chromosomal distribution of all detected significantly differential expressed genes between white and yellow genotypes across all 18 *Manihot esculenta* chromosomes. Black bars at the bottom depict the localization of genes on the respective chromosome. The line in the background is representing the number of transcripts for that position. Green dots are referring to the significant differentially expressed genes for that chromosome. Y- axis and equivalent height of green dots is referring to changes in gene expression comparing white with yellow genotypes – positive values referring to a higher expression in white, whereas negative values representing a higher expression in yellow genotypes.

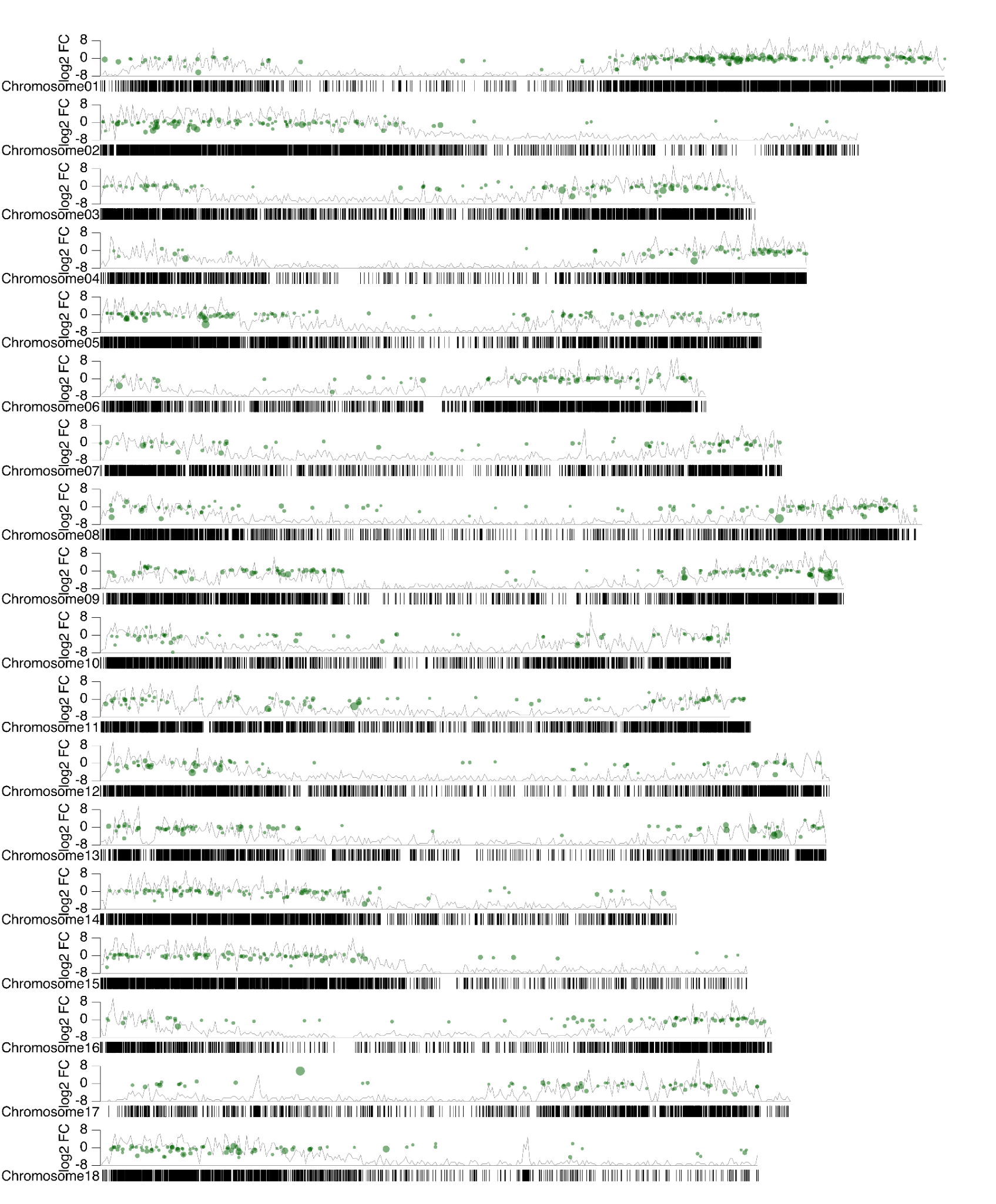

**Supplementary material - Figure S2**

Functional enrichment analysis for genes showing significant correlation to starch after correcting for carotenoid content as covariate. The upper left part is depicting KEGG categorization of transcripts positively correlated to starch in white genotypes, and the right upper part is showing the KEGG categorization of negatively correlated transcripts. The lower part represents the KEGG categorization for the transcript abundance in yellow genotypes positively (left) and negatively (right) correlated to starch.
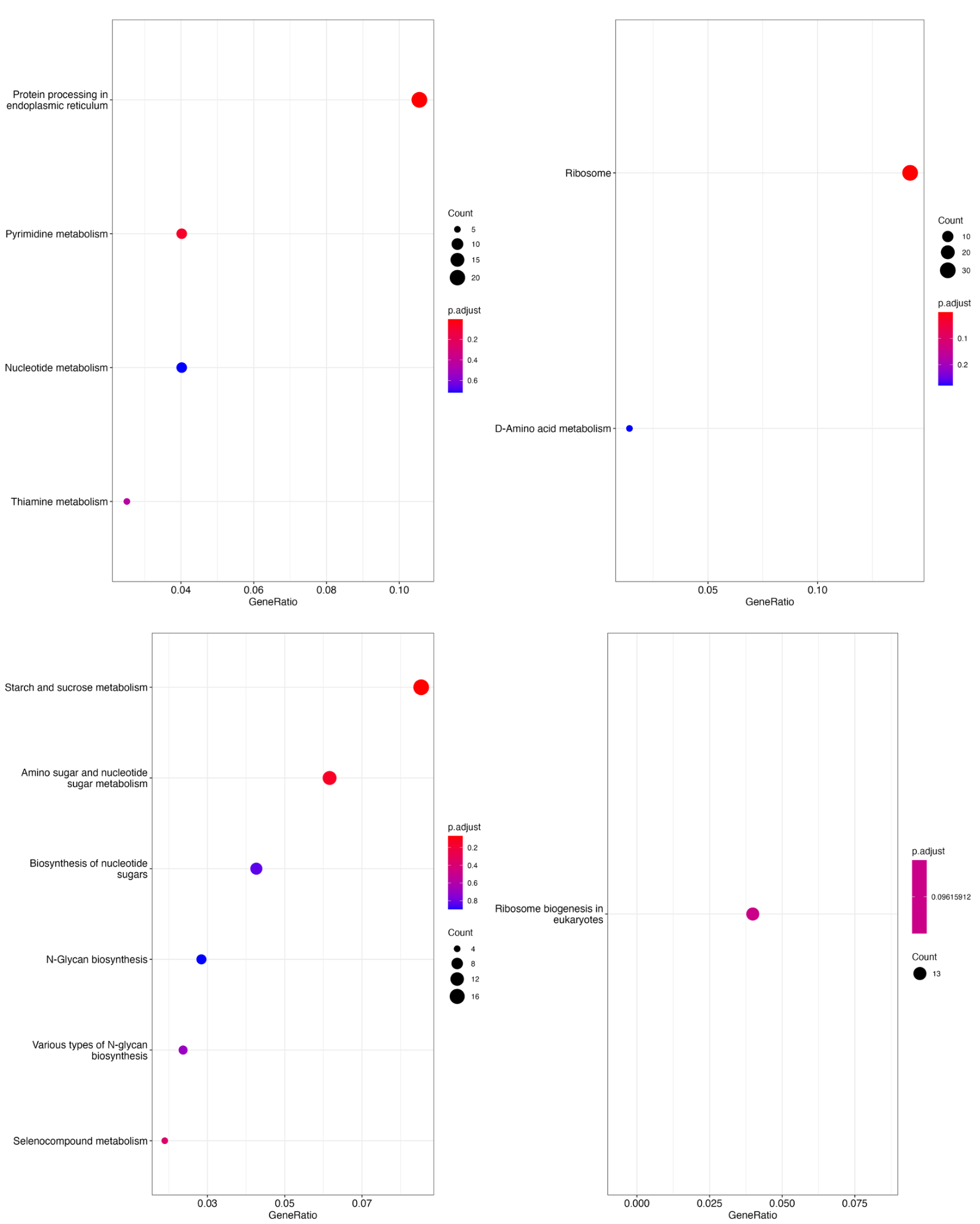

**R Session Info**

R version 4.1.0 (2021-05-18)

Platform: x86_64-apple-darwin17.0 (64-bit)

Running under: macOS Big Sur 10.16

Matrix products: default

BLAS: /Library/Frameworks/R.framework/Versions/4.1/Resources/lib/libRblas.dylib

LAPACK: /Library/Frameworks/R.framework/Versions/4.1/Resources/lib/libRlapack.dylib

locale:

[1] en_US.UTF-8/en_US.UTF-8/en_US.UTF-8/C/en_US.UTF-8/en_US.UTF-8

attached base packages:

[1] stats4	stats	graphics grDevices utils	datasets methods	base

other attached packages:

[1] car_3.1-2	carData_3.0-5	lme4_1.1-34	karyoploteR_1.18.0

[5] yarrr_0.1.5	circlize_0.4.15	BayesFactor_0.9.12-4.4	Matrix_1.5-1

[9] coda_0.19-4	jpeg_0.1-10	regioneR_1.24.0	GenomicFeatures_1.44.2

[13] AnnotationDbi_1.54.1	corrplot_0.92	reshape2_1.4.4	EnhancedVolcano_1.10.0

[17] ggrepel_0.9.3	factoextra_1.0.7	cowplot_1.1.1	ggpubr_0.6.0

[21] ppcor_1.1	MASS_7.3-60	lubridate_1.9.2	forcats_1.0.0

[25] stringr_1.5.0	purrr_1.0.2	readr_2.1.4	tidyr_1.3.0

[29] tidyverse_2.0.0	ggplot2_3.4.3	clusterProfiler_4.0.5	DESeq2_1.32.0

[33] SummarizedExperiment_1.22.0 Biobase_2.52.0	MatrixGenerics_1.6.0	matrixStats_1.0.0

[37] GenomicRanges_1.44.0	GenomeInfoDb_1.28.4	IRanges_2.28.0	S4Vectors_0.32.4

[41] BiocGenerics_0.40.0	tibble_3.2.1	dplyr_1.1.2

loaded via a namespace (and not attached):

[1] rappdirs_0.3.3	rtracklayer_1.52.1	bamsignals_1.24.0	ragg_1.2.5

[5] bezier_1.1.2	bit64_4.0.5	knitr_1.43	DelayedArray_0.20.0

[9] data.table_1.14.8	rpart_4.1.19	KEGGREST_1.32.0	RCurl_1.98-1.12

[13] AnnotationFilter_1.16.0 generics_0.1.3	RSQLite_2.3.1	shadowtext_0.1.2

[17] bit_4.0.5	tzdb_0.4.0	enrichplot_1.12.3	xml2_1.3.5

[21] viridis_0.6.4	xfun_0.40	hms_1.1.3	evaluate_0.21

[25] fansi_1.0.4	restfulr_0.0.15	progress_1.2.2	dbplyr_2.3.3

[29] igraph_1.4.0	DBI_1.1.3	geneplotter_1.70.0	htmlwidgets_1.6.2

[33] backports_1.4.1	annotate_1.70.0	biomaRt_2.48.3	vctrs_0.6.3

[37] ensembldb_2.16.4	abind_1.4-5	cachem_1.0.8	withr_2.5.0

[41] ggforce_0.4.1	BSgenome_1.60.0	checkmate_2.2.0	GenomicAlignments_1.28.0

[45] treeio_1.16.2	prettyunits_1.1.1	cluster_2.1.4	DOSE_3.18.3

[49] ape_5.7-1	lazyeval_0.2.2	crayon_1.5.2	genefilter_1.74.1

[53] labeling_0.4.2	pkgconfig_2.0.3	tweenr_2.0.2	nlme_3.1-162

[57] vipor_0.4.5	ProtGenerics_1.24.0	nnet_7.3-19	rlang_1.1.1

[61] lifecycle_1.0.3	MatrixModels_0.5-1	downloader_0.4	filelock_1.0.2

[65] extrafontdb_1.0	BiocFileCache_2.0.0	dichromat_2.0-0.1	ggrastr_1.0.2

[69] polyclip_1.10-4	aplot_0.2.0	boot_1.3-28.1	base64enc_0.1-3

[73] beeswarm_0.4.0	GlobalOptions_0.1.2	png_0.1-8	viridisLite_0.4.2

[77] rjson_0.2.21	bitops_1.0-7	KernSmooth_2.23-20	Biostrings_2.60.2

[81] blob_1.2.4	shape_1.4.6	qvalue_2.24.0	rstatix_0.7.2

[85] gridGraphics_0.5-1	ggsignif_0.6.4	scales_1.2.1	memoise_2.0.1

[89] magrittr_2.0.3	plyr_1.8.8	zlibbioc_1.38.0	compiler_4.1.0

[93] scatterpie_0.2.1	BiocIO_1.2.0	RColorBrewer_1.1-3	ash_1.0-15

[97] Rsamtools_2.8.0	cli_3.6.1	XVector_0.32.0	patchwork_1.1.3

[101] pbapply_1.7-2	htmlTable_2.4.1	Formula_1.2-5	mgcv_1.8-41

[105] tidyselect_1.2.0	stringi_1.7.12	textshaping_0.3.6	proj4_1.0-12

[109] yaml_2.3.7	GOSemSim_2.18.1	locfit_1.5-9.8	grid_4.1.0

[113] VariantAnnotation_1.38.0 fastmatch_1.1-4	tools_4.1.0	timechange_0.2.0

[117] parallel_4.1.0	rstudioapi_0.15.0	foreign_0.8-84	gridExtra_2.3

[121] farver_2.1.1	ggraph_2.1.0	digest_0.6.33	Rcpp_1.0.11

[125] broom_1.0.5	ggalt_0.4.0	httr_1.4.7	biovizBase_1.40.0

[129] colorspace_2.1-0	XML_3.99-0.14	splines_4.1.0	yulab.utils_0.0.7

[133] tidytree_0.4.5	graphlayouts_1.0.0	ggplotify_0.1.2	systemfonts_1.0.4

[137] xtable_1.8-4	nloptr_2.0.3	jsonlite_1.8.7	ggtree_3.0.4

[141] tidygraph_1.2.3	ggfun_0.1.2	R6_2.5.1	Hmisc_5.1-0

[145] pillar_1.9.0	htmltools_0.5.6	glue_1.6.2	fastmap_1.1.1

[149] minqa_1.2.5	BiocParallel_1.28.3	maps_3.4.1	fgsea_1.18.0

[153] mvtnorm_1.1-3	utf8_1.2.3	lattice_0.21-8	curl_5.0.2

[157] ggbeeswarm_0.7.2	GO.db_3.13.0	Rttf2pt1_1.3.12	survival_3.5-7

[161] rmarkdown_2.24	munsell_0.5.0	DO.db_2.9	GenomeInfoDbData_1.2.6

[165] gtable_0.3.4	extrafont_0.19
